# Conserved *Cis*-Acting Range Extender Element Mediates Extreme Long-Range Enhancer Activity in Mammals

**DOI:** 10.1101/2024.05.26.595809

**Authors:** Grace Bower, Ethan W. Hollingsworth, Sandra Jacinto, Benjamin Clock, Kaitlyn Cao, Mandy Liu, Adam Dziulko, Ana Alcaina-Caro, Qianlan Xu, Dorota Skowronska-Krawczyk, Javier Lopez-Rios, Diane E. Dickel, Anaïs F. Bardet, Len A. Pennacchio, Axel Visel, Evgeny Z. Kvon

**Affiliations:** Department of Developmental and Cell Biology, University of California, Irvine, CA 92967, USA; Medical Scientist Training Program, University of California, Irvine, CA 92967, USA; Environmental Genomics and Systems Biology Division, Lawrence Berkeley National Laboratory, Berkeley, CA 94720, USA; Centro Andaluz de Biología del Desarrollo (CABD), CSIC-Universidad Pablo de Olavide-Junta de Andalucía, Seville, 41013, Spain; Department of Physiology and Biophysics, Department of Ophthalmology, Center for Translational Vision Research, School of Medicine, University of California, Irvine, CA, USA; Institut de Génétique et de Biologie Moléculaire et Cellulaire (IGBMC), Université de Strasbourg, CNRS UMR7104, INSERM U1258, 67400 Illkirch, France; U.S. Department of Energy Joint Genome Institute, Walnut Creek, CA 94598, USA; Comparative Biochemistry Program, University of California, Berkeley, CA 94720, USA; School of Natural Sciences, University of California, Merced, CA 95343, USA

## Abstract

While most mammalian enhancers regulate their cognate promoters over moderate distances of tens of kilobases (kb), some enhancers act over distances in the megabase range. The sequence features enabling such extreme-distance enhancer-promoter interactions remain elusive. Here, we used *in vivo* enhancer replacement experiments in mice to show that short– and medium-range enhancers cannot initiate gene expression at extreme-distance range. We uncover a novel conserved *cis*-acting element, <u>R</u>ange <u>EX</u>tender (REX), that confers extreme-distance regulatory activity and is located next to a long-range enhancer of *Sall1*. The REX element itself has no endogenous enhancer activity. However, addition of the REX to other short– and mid-range enhancers substantially increases their genomic interaction range. In the most extreme example observed, addition of the REX increased the range of an enhancer by an order of magnitude, from its native 71kb to 840kb. The REX element contains highly conserved [C/T]AATTA homeodomain motifs. These motifs are enriched around long-range limb enhancers genome-wide, including the ZRS, a benchmark long-range limb enhancer of *Shh*. Mutating the [C/T]AATTA motifs within the ZRS does not affect its limb-specific enhancer activity at short range, but selectively abolishes its long-range activity, resulting in severe limb reduction in knock-in mice. In summary, we identify a sequence signature globally associated with long-range enhancer-promoter interactions and describe a prototypical REX element that is necessary and sufficient to confer extreme-distance gene activation by remote enhancers.

## Introduction

Transcriptional enhancers are abundant *cis*-acting non-coding genomic elements that activate gene expression across different cell types of the organism in response to internal and external signals (Long et al., 2016; Serebreni & Stark, 2021; Shlyueva et al., 2014; Spitz & Furlong, 2012). Enhancers often regulate their target promoters across long genomic distances (Benabdallah et al., 2019; Z. Chen et al., 2024; Lettice et al., 2002; Long et al., 2020; Padhi et al., 2021; Robson et al., 2019; Williamson et al., 2016; Yanchus et al., 2022). Disruption of this long-range gene regulation often leads to diseases ranging from developmental anomalies to cancer (Franke et al., 2016; Long et al., 2020; Lupiáñez et al., 2015; Northcott et al., 2014; Wang et al., 2021; Weischenfeldt & Ibrahim, 2023; Zaugg et al., 2022).

A central question in gene regulation is how remote enhancers precisely relay regulatory information to their target promoters located hundreds of thousands or even millions of base pairs away, often without impacting the expression of intervening genes. While enhancers come into physical proximity to activate target promoters, the mechanisms mediating these functionally important interactions are only partially understood (Benabdallah et al., 2019; Z. Chen et al., 2024; Hafner & Boettiger, 2023; Robson et al., 2019; Williamson et al., 2016). Mounting evidence indicates that higher-order 3D-chromatin organization and structural CTCF/cohesin proteins support enhancer-promoter (E-P) interactions by restricting E–P communication to so-called topologically associating domains (TADs) (Dixon et al., 2012; Kane et al., 2022; Nora et al., 2012; Rinzema et al., 2022; Robson et al., 2019; Schoenfelder & Fraser, 2019). However, global disruption of higher-order 3D-chromatin organization and loop extrusion in CTCF and cohesin-depleted cells only partially disrupts E–P communication and gene expression (Hsieh et al., 2022; Nora et al., 2017; Rao et al., 2017; Schwarzer et al., 2017). Furthermore, developmental gene expression is also surprisingly robust to CTCF binding site deletions and structural perturbations affecting TADs (Barutcu et al., 2018; Despang et al., 2019; Ghavi-Helm et al., 2019; Paliou et al., 2019; Seitan et al., 2013; Soshnikova et al., 2010; Ushiki et al., 2021; Williamson et al., 2019). Therefore, which additional genetic factors establish and maintain long-range E–P communication during mammalian development remains poorly understood (Furlong & Levine, 2018; Hafner & Boettiger, 2023; Zaugg et al., 2022). Identifying such factors is crucial for achieving an in-depth understanding of developmental processes and delineating disease mechanisms linked to disruption of long-range gene regulation.

In the present study, we uncovered a unique sequence signature comprised of [C/T]AATTA homeodomain (HD) motifs that mediates long-range enhancer–promoter communication in developing limb buds. Deletion of these motifs selectively abolishes distal enhancer activity but not its short-range ability. The [C/T]AATTA motifs are enriched at long-distance limb enhancer loci genome-wide and their presence correlates with enhancer–promoter distance. We also characterize a remarkable and extreme case in which the [C/T]AATTA motifs are located spatially separate from the enhancer within a genomic element which we termed the <u>R</u>ange <u>EX</u>tender (REX). The addition of the REX element can convert a short-range enhancer into a long-range enhancer that can act over 840 kb of genomic space. Our results indicate that short– or medium-range enhancers cannot function at extreme-distance range and establish that spatial specificity and long-distance activity are two separate and separable aspects of enhancer function.

## Results

### Short– and medium-range limb enhancers cannot act over long genomic distances

To assess the potential of short– and medium-range enhancers to act over long genomic distances, we performed a series of enhancer replacement experiments at the *Shh* genomic locus. Limb-specific expression of *Shh* is controlled by the ZRS (zone of polarizing activity (ZPA) regulatory sequence, also known as MFCS1), located at an extreme distance of ∼850 kb from its target promoter in mice (Lettice et al., 2003). Mice deficient for the endogenous 1.3-kb ZRS region fail to initiate any *Shh* expression in developing limb buds and display truncated limb phenotypes indistinguishable from complete loss of SHH function in the limb (Sagai et al., 2005). Because of the clear phenotypic readout and lack of redundancy, ZRS-*Shh* is an ideal locus for assessing the long-range enhancer activity of transplanted enhancers.

To perform enhancer replacement experiments, we selected four previously characterized developmental limb bud enhancers from other genomic loci based on their ability to drive robust LacZ reporter expression in the developing limb bud mesenchyme (MM1492 and MM1564 (from mouse), and HS1516 and HS72 (from human); **Fig. 1A**,**B**) (Dickel et al., 2013; Lettice et al., 2003; Osterwalder et al., 2014; Visel et al., 2007). The HS72, HS1516, and MM1492 enhancers also drive LacZ reporter expression in the ZPA, a posterior domain of the limb mesenchyme where ZRS activates *Shh* gene expression (**Fig. 1B**). All four enhancers are evolutionarily conserved and marked by H3K27ac in limb buds, a histone modification associated with active enhancers (**Fig. 1A**). To determine the genomic distance at which these enhancers act on their native promoters and to ensure that they are accessible to transcription factors (TFs) in the same cell types as the ZRS, we performed single nuclei ATAC-seq and RNA-seq multiome profiling of 14,000 cells from an E11.5 hindlimb bud (see **Methods**). After clustering all 14,000 cells based on their chromatin and gene expression profiles, we annotated sixteen clusters representing all major cell types in the developing limb bud (**Fig. S1, Table S1,** and **Methods**). The HS72, HS1516, and MM1492 enhancers display strong DNA accessibility in *Shh* positive ZPA cells, consistent with their ability to drive robust reporter expression in posterior limb mesenchyme in transgenic mouse embryos (**Fig. 1A** and **S1C**).

**Figure 1:**
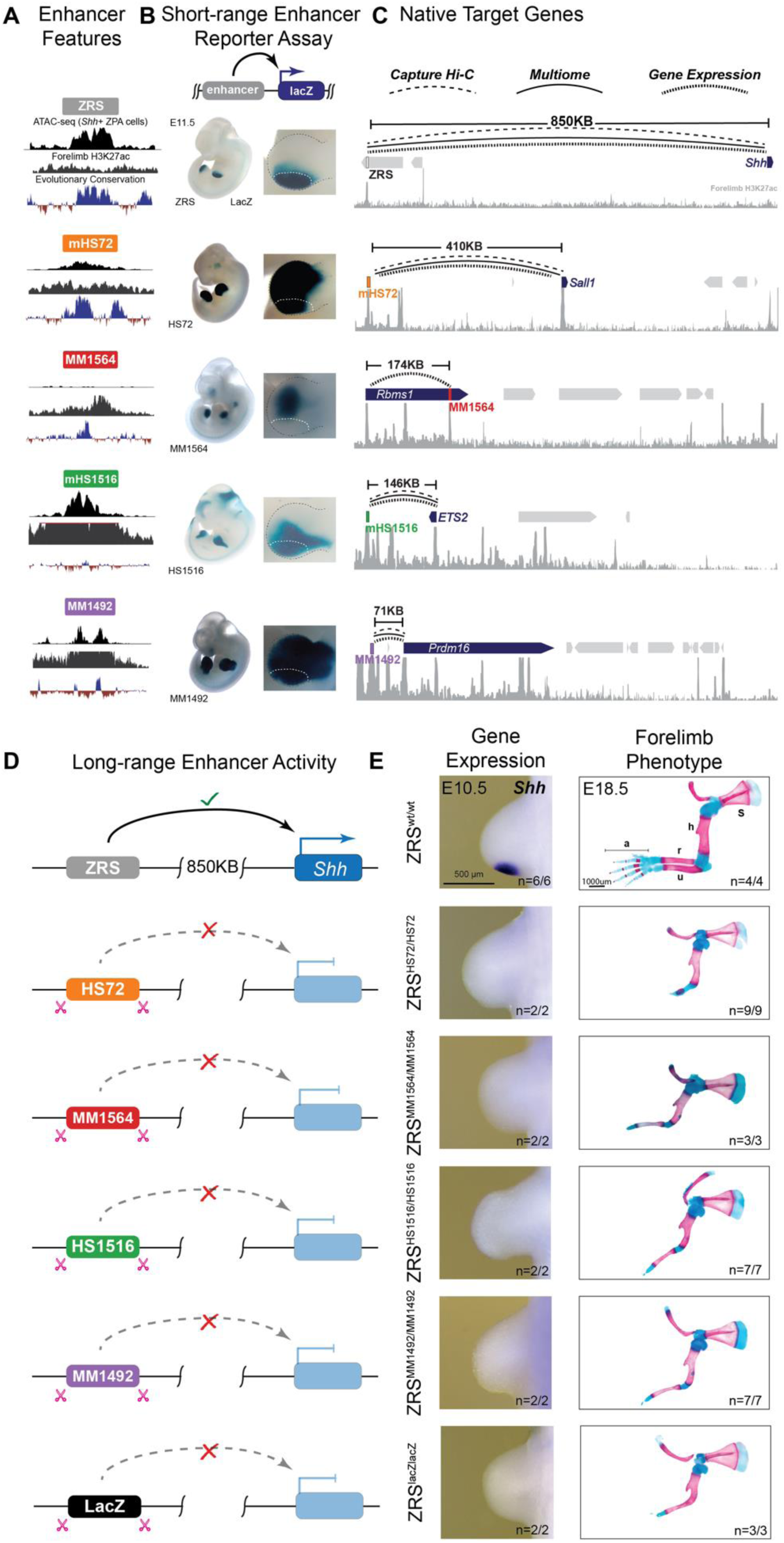
Transplanted Enhancers Lack Long-Range Limb Activity in Knock-in Mice. (**A**) All selected limb enhancers (colored blocks) are marked by H3K27 acetylation and open chromatin and contain a conserved core sequence. For each enhancer, E11.5 forelimb H3K27ac ChIP-seq (Lex et al., 2020)), pseudobulk *Shh*+ ZPA scATAC-seq cluster and placental conservation tracks are shown. mHS72 and mHS1516 are mouse orthologous of HS72 and HS1516. **(B)** Corresponding enhancer activities in transgenic E11.5 mouse embryos. Zoomed in forelimb buds are shown on the right. The white dotted line demarcates the approximate ZPA region. (**C**) Corresponding genomic regions where curved lines indicate putative E– P interactions supported by capture Hi-C (Z. Chen et al., 2024), multiome analysis of E11.5 mouse limb buds (scATAC-seq + scRNA-seq correlation; this study) or matched enhancer activity and gene expression in limb buds of E11.5 or E10.5 embryos (**Fig. S2D**). H3K27ac ChIP-seq signal from E11.5 forelimbs is shown underneath each region (gray). **(D-E**) Limb phenotyping of knock-in mice with transplanted enhancers. The ZRS is located ∼850 kb away from the promoter of *Shh*. (**D**) Schematic mouse *Shh* loci with the ZRS replaced by HS72, MM1564, HS1516, and MM1492 limb enhancers or a fragment of the *lacZ* sequence are shown. (**E**) Comparative *Shh* mRNA in situ hybridization analysis in wild type and knock-in mouse embryos during forelimb bud development (first column). The genotypes of the embryos are shown on the left. Corresponding skeletal preparations of E18.5 mouse embryos are shown in the second column: s, scapula; h, humerus; r, radius; u, ulna; a, autopod. The number of embryos that exhibited the representative limb phenotype over the total number of embryos with the genotype is indicated. See **Fig. S2** for more details, including hindlimb bud analysis. The coordinates of enhancers used in B, D and E are: ZRS (chr5:29,314,497-29,315,844; mm10), HS72 (chr16:51,623,899-51,624,805; hg38), MM1564 (chr2:60,785,660-60,787,563; mm10), HS1516 (hg38_chr21:38,989,433-38,991,200; hg38) and MM1492 (chr4:154,707,415-154,711,162; mm10).

To link limb enhancers to their putative target genes, we used the correlation between gene expression and open chromatin peaks from multiome profiling and previously published E–P physical interaction data based on tissue-resolved high-resolution enhancer-capture-Hi-C experiments (Z. Chen et al., 2024) (See **Methods**). We identified *Prdm16* as the target for MM1492 (71 kb away), *ETS2* as the target for mHS1516 (150 kb away), and *Sall1* as the target for mHS72 (407 kb away) (**Fig. 1C**) (Dickel et al., 2013; Osterwalder et al., 2018). We then manually matched enhancer activity and the expression patterns of genes located within the same TADs in E11.5 mouse embryos. This analysis confirmed the above E–P links and additionally identified *Rbms1* as a putative target for MM1564 (174 kb away; **Fig. 1B**,**C** and **Fig. S2C**).

We next employed genome editing to create a series of knock-in (KI) mice in which the functionally critical 1.3-kb region of the ZRS (**Fig. 1D** and **Fig. S2**) was replaced with selected limb bud enhancers from other genomic loci. Surprisingly, replacing the mouse ZRS with HS72, MM1564, HS1516, or MM1492 limb enhancers resulted in a loss of detectable *Shh* expression in the limb bud (**Fig. 1E**). To assess the extent to which replacing the ZRS with a heterologous limb enhancer affects *in vivo* development we examined skeletal morphology in E18.5 knock-in mice. Consistent with the loss of limb-specific *Shh* expression, all four knock-in mouse strains displayed a truncated limb phenotype affecting both the fore– and hindlimbs. The limb phenotypes were indistinguishable from the phenotype caused by replacement of the ZRS with a nonfunctional control sequence of similar length (**Fig. 1E** and **Fig. S2D**). These results indicate that despite the presence of *bona fide* enhancer features capable of driving strong limb gene expression in the context of a transgene, all four enhancers lack long-range activity and cannot support *Shh*-mediated limb development. These results also suggest that developmental enhancers with identical tissue specificity in short-range transgenic reporter assays are not interchangeable within the context of the genome due to differences in their ability to act over long distances.

### Transplanted enhancers maintain their endogenous DNA accessibility pattern

It is possible that silencing of the inserted region or failure of the insertions to create an open chromatin environment explains the inability of transplanted regions to act as long-range enhancers (Haruyama et al., 2009; Martin & Whitelaw, 1996). To test whether enhancers maintain their endogenous chromatin architecture at a remote ectopic location, we performed ATAC-seq experiments in ZPA region of fore– and hindlimb buds of E11.5 mice heterozygous for the ZRS^HS72^ and ZRS^HS1516^ knock-in alleles.

HS72 and HS1516 are human enhancers that are highly conserved in mice but contain substitutions that allow discrimination of ATAC-seq reads from transplanted human and orthologous endogenous mouse enhancers (**Fig. S3** and **Methods**). ZRS^HS72^ and ZRS^HS1516^ heterozygous mice formed normal limbs, which allowed us to directly compare chromatin accessibility at the transplanted enhancer allele and the wild-type ZRS enhancer allele in fully developed limb bud tissue from the same mouse. Allele-specific ATAC-seq profiles demonstrated that both transplanted enhancers and orthologous mouse enhancers had an accessible open chromatin architecture (**Fig. 2A**). These results indicate that the heterologous HS72 and HS1516 limb enhancers were accessible when transplanted yet were unable to activate gene expression remotely, further supporting that their inability to activate the *Shh* promoter in ZRS replacement experiments is due to limitations in their genomic interaction range.

**Figure 2:**
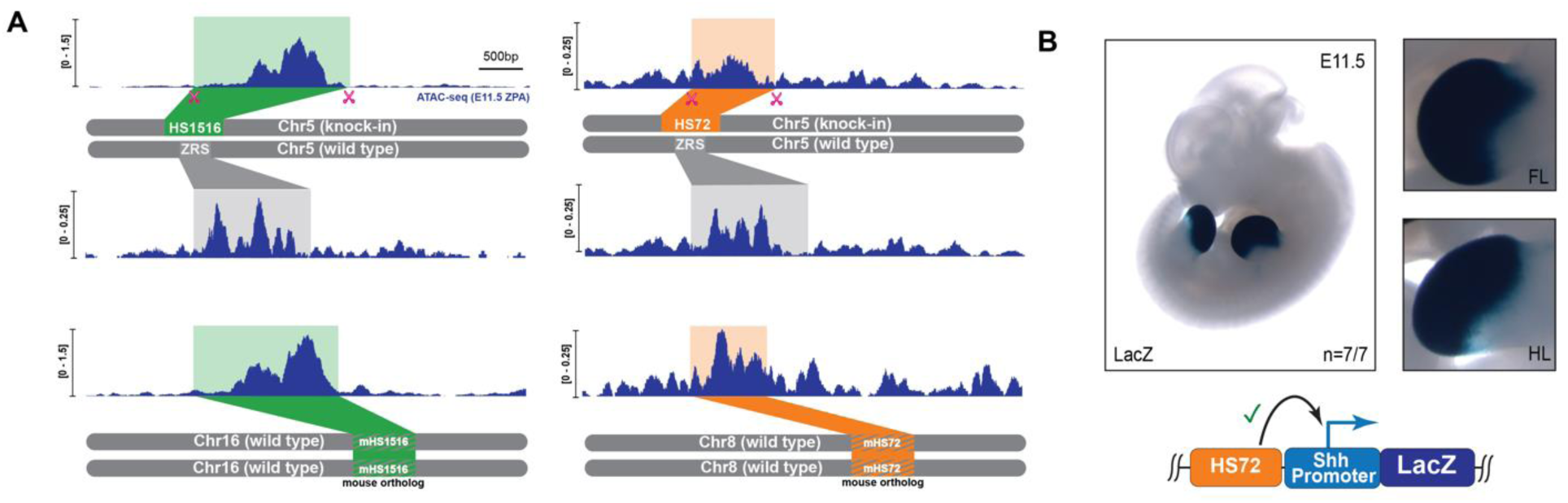
Transplanted enhancers maintain open chromatin structure at the knock-in site and drive functional SHH expression in the limb when proximal to the *Shh* promoter. (**A**) Top: allele-specific ZPA ATAC-seq signal at the transplanted HS1516 (green box) or HS72 (orange box) enhancers and the corresponding wild-type ZRS locus. Bottom: ATAC-seq profiles at endogenous mHS1516 (left; chr16:95,847,911-95,849,564) and mHS72 (right; chr8:89,454,508-89,455,383) mouse orthologous enhancers (green and orange striped boxes). (**B**) LacZ-stained transgenic E11.5 mouse embryo carrying an HS72 limb enhancer inserted upstream of the *Shh* promoter (light blue) and *lacZ* reporter gene (dark blue). The number of embryos with robust LacZ staining in limb buds over the total number of transgenic embryos screened is indicated.

### Heterologous limb enhancers can activate *Shh* expression in a transgene

The inability of heterologous limb enhancers to activate *Shh* expression was surprising, given their robust enhancer features and ability to establish open chromatin at the transplanted location. Previously, all four limb enhancers were characterized in transgenic mice using a well-established *hsp68* minimal promoter (Kothary et al., 1989; Pennacchio et al., 2006), but their compatibility, or lack thereof, with the *Shh* promoter is unknown. To test if E–P incompatibility (Butler & Kadonaga, 2001; Juven-Gershon & Kadonaga, 2010; Zabidi et al., 2015) could play a role in the observed differences in gene activation, we placed the HS72 enhancer upstream of the mouse *Shh* promoter and the *LacZ* reporter gene and injected the resulting construct into fertilized mouse eggs. All transgenic mouse embryos (7/7) displayed strong *LacZ* expression in the limb buds, which was identical to HS72’s activity with the *hsp68* promoter, indicating that the HS72 enhancer is fully capable of activating the *Shh* promoter (**Fig. 1A** and **Fig. 2B**). Moreover, we previously developed a site-directed reporter system and showed that 32 enhancers maintained their *in vivo* activity whether driving a minimal *hsp68* or a new minimal *Shh* promoter (Kvon et al., 2020; Osterwalder et al., 2022; Snetkova et al., 2021). We also placed the HS72 enhancer upstream of the *Shh* promoter and the *Shh* coding sequence and integrated this transgene at the safe-harbor location of the mouse genome (Kvon et al., 2020). A typical readout of ectopic *Shh* expression is the preaxial polydactyly observed when *Shh* is expressed beyond the ZPA in the limb bud (Lettice et al., 2003, 2002). Since HS72 is active more broadly than the ZRS in limb mesenchyme, a polydactyly phenotype would indicate functional SHH signaling. Indeed, all transgenic mouse embryos (6/6) developed strong polydactyly in both fore– and hindlimbs, consistent with the HS72 enhancer driving broad functional *Shh* activity (Lettice et al., 2008) (**Fig. S4D**). These results indicate that intrinsic E–P specificity is likely not the reason for observed differences between short– and long-range activation.

### A range extender (REX) element is required for long-range gene activation by a *Sall1* enhancer

All four heterologous limb enhancers lie closer to their putative target genes than the ZRS (∼850 kb), suggesting that their failure to activate *Shh* in genomic replacement experiments is due to the extreme distance from the target promoter (**Fig. 1C**). To explore possible factors in long-distance activation, we focused on the heterologous enhancer with the longest native activity range, HS72, which is located ∼407 kb away from the *Sall1* promoter (**Fig. 1C**). We hypothesized that there may be additional functional sequences associated with distant-acting enhancers that support long-range activation. We identified a highly conserved block of sequence located adjacent to the HS72 enhancer (**Fig. 3A**). This sequence is not required to drive limb-specific activity in reporter assays (**Figs. 1B**, **2B**, and **S4D**), but because of its strong conservation and position, we hypothesized that this sequence might support enhancer activity, including the ability to activate gene expression over remote distances. To test this hypothesis, we generated knock-in mice in which the ZRS was replaced with an extended version of the HS72 enhancer that included this highly conserved downstream sequence (**Fig. 3B**).

**Figure 3:**
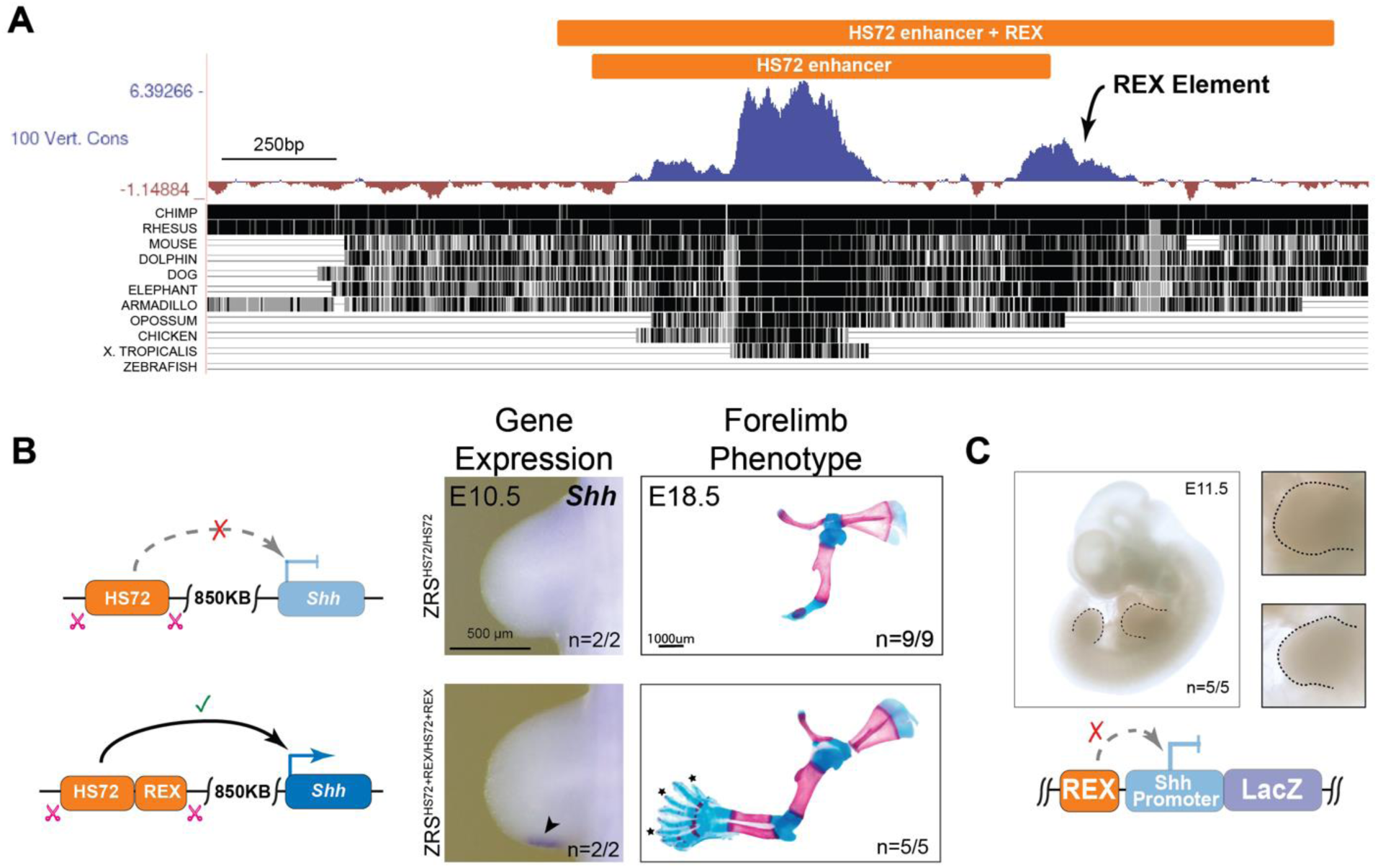
The REX element is necessary for long-range activation of *Shh* by a heterologous enhancer. (**A**) An evolutionary conserved element of unknown function is located adjacent to the human HS72 enhancer. The HS72 enhancer region is shown together with evolutionary conservation tracks. (**B**) Replacement of the ZRS with an extended version of the HS72 enhancer (chr16:51,623,658-51,625,572; hg38) containing the REX element results in initiation of *Shh* expression in developing limb buds and full limb outgrowth in knock-in mice. *Extra digits (polydactyly) in ZRS^HS72+REX^/ZRS^HS72+REX^ embryos. See **Fig. S4** for hindlimb and E11.5 *Shh* expression analysis. (**C**) The REX element lacks classical enhancer activity in E11.5 transgenic embryos when placed upstream of the *Shh* promoter and lacZ reporter gene. The number of transgenic embryos with no LacZ activity in the limb over the total number of transgenic embryos screened is indicated.

Consistent with our prediction, addition of this uncharacterized sequence which we termed the “**<u>R</u>**ange **<u>Ex</u>**tender (<u>REX</u>) element” to the transplanted HS72 enhancer was sufficient to initiate *Shh* expression in the limb and resulted in full limb outgrowth with formation of all distal limb elements including fully formed zeugopod and autopod in ZRS^HS72REX/HS72REX^ mice (**Fig. 3B** and **Fig. S4A**). ZRS^HS72REX/HS72REX^ mice also displayed polydactyly on both fore– and hindlimbs, consistent with ectopic *Shh* expression in limb buds driven by broader HS72 enhancer activity (**Figs. 1B**, **2B**, **3B**, and **S4**).

Notably, the REX element alone does not act as a classical enhancer. When placed upstream of the minimal promoter in a transgene, it is unable to drive *LacZ* reporter expression in transgenic mouse embryos (**Fig. 3C**). Taken together, these results indicate that the REX element does not have enhancer activity on its own but is required for long-range heterologous enhancer activation at the *Shh* locus.

### REX confers long-distance activation range to a heterologous short-range enhancer

To test if the REX element can extend the range of heterologous short-range enhancers, we generated a knock-in mouse line where the ZRS was replaced with a chimeric element consisting of the shortest-range heterologous limb enhancer from our test set (MM1492, 71 kb native E—P range) appended by the REX element (MM1492REX; **Fig. S5A**). Knock-in of this chimeric element was sufficient to induce *Shh* expression in the limb bud and resulted in a fully developed zeugopod and autopod in the ZRS^MM1492REX/MM1492REX^ mice (**Fig. 4** and **Fig. S5B**). ZRS^MM1492REX/MM1492REX^ mice also displayed polydactyly, consistent with broad limb activity of MM1492. Taken together, these experiments show that combining a short-range enhancer with the REX element and placing them at an ectopic remote location is sufficient to induce long-range gene activation. These results also suggest that the REX element can act in a modular fashion to facilitate long-range enhancer action.

**Figure 4:**
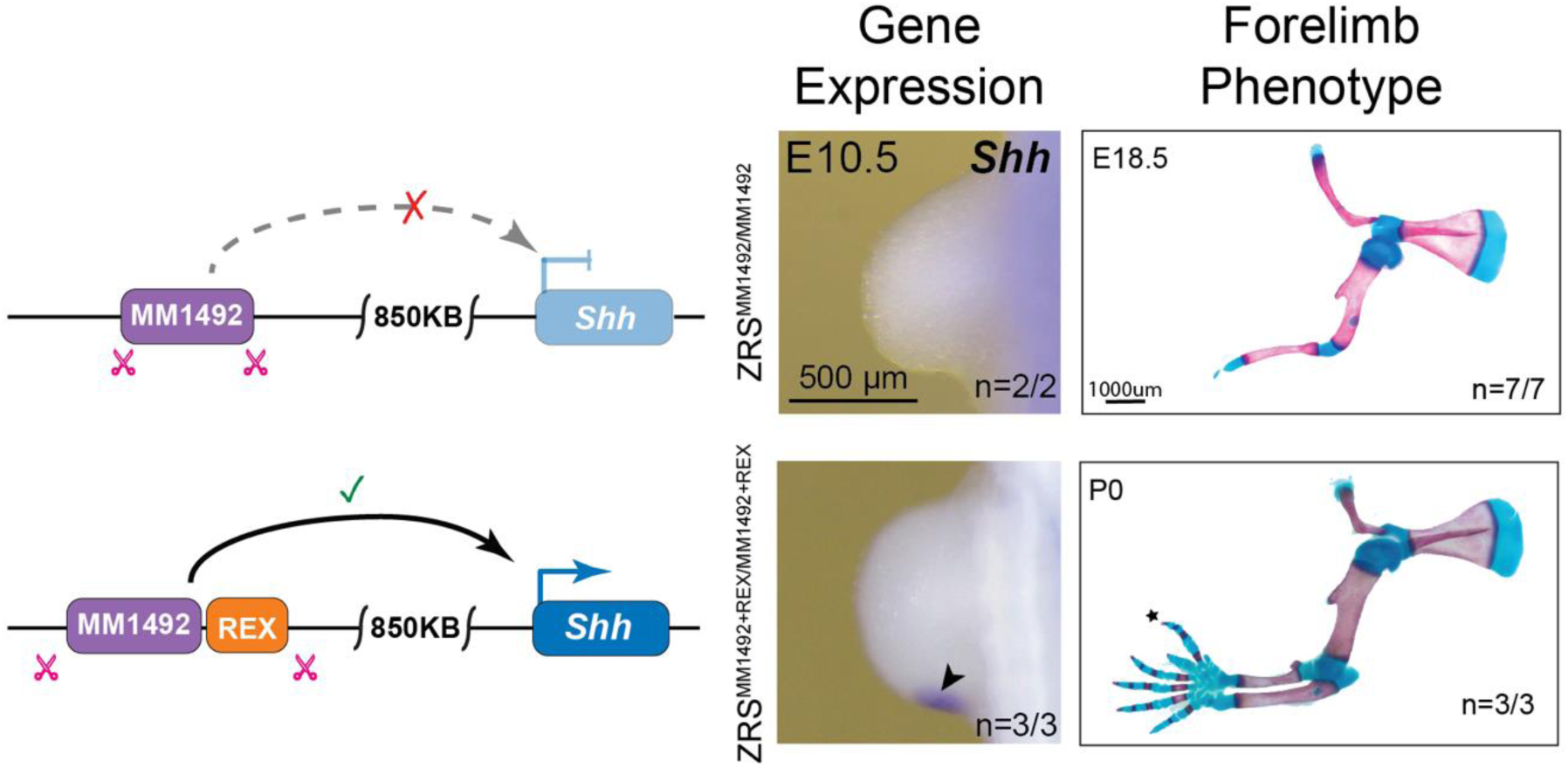
The REX element can convert a short-range enhancer into a long-range enhancer. Replacement of the ZRS with a chimeric cis-regulatory element consisting of the short-range MM1492 enhancer and the REX element from the HS72 enhancer region results in the initiation of *Shh* expression in developing limb buds at E10.5 and full limb outgrowth at P0 in knock-in mice. *Extra digits (polydactyly) in ZRS^MM1492REX^/ZRS^MM1492REX^ embryos. See **Fig. S5B-C** for hindlimb and E10.5 *Shh* expression analysis.

### REX element contains [C/T]AATTA homeodomain motifs that are globally linked to extreme long-range activation

To identify specific TFs that may be involved in REX-mediated long-range enhancer action, we examined potential transcription factor binding sites within the REX element. We scanned the REX element sequence for candidate TF binding sites and identified conserved motifs whose sequences matched binding preferences for LHX2 and LHX9 homeodomain (HD) TFs as well as LEF1 (**Figs. 5A**). LHX2 and LEF1 motifs are evolutionarily conserved across all placental mammals, and the LHX9 motif is conserved across eutherian mammals but not in marsupials (**Figs. 5A** and **S6A)**.

**Figure 5:**
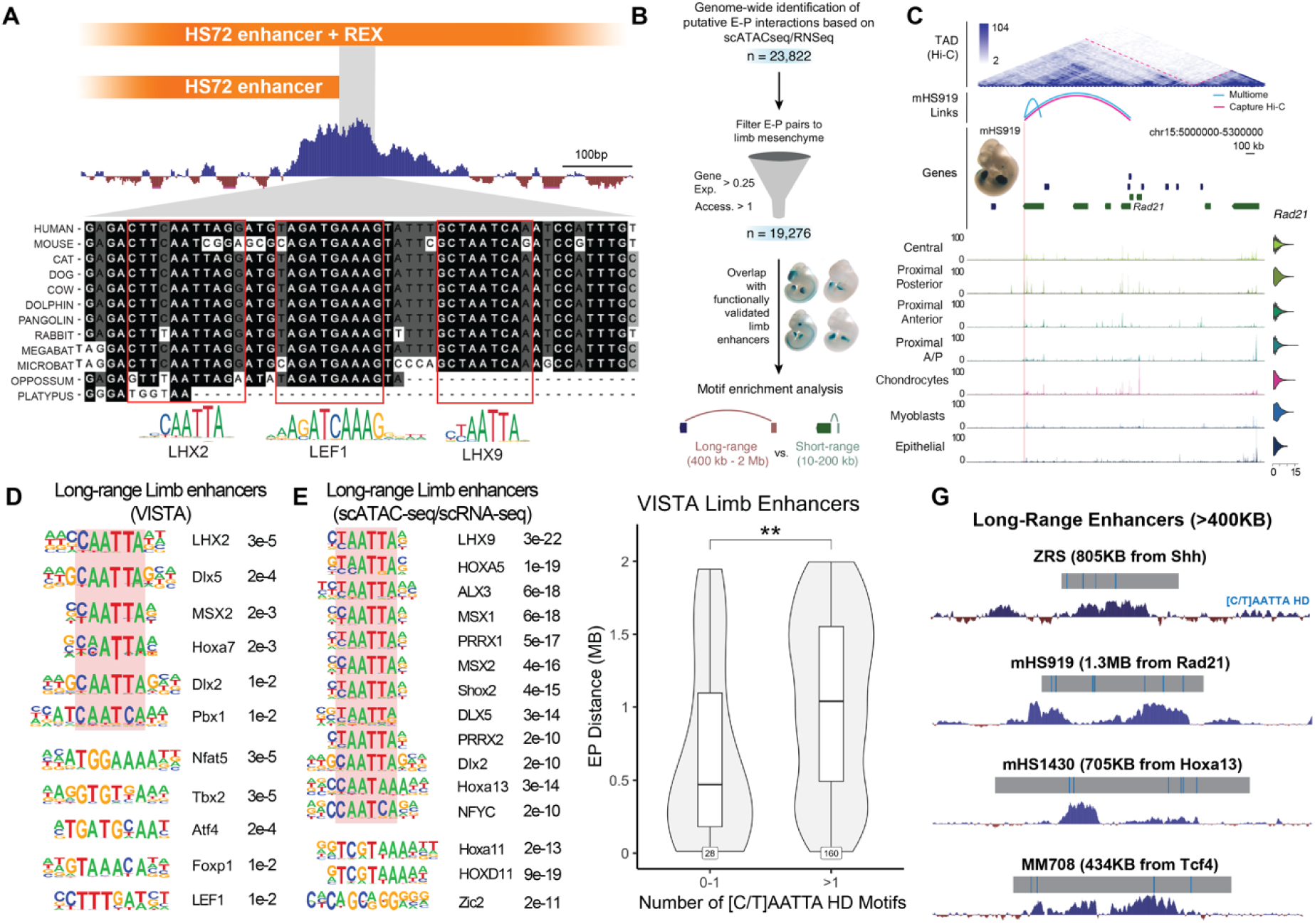
[C/T]AATTA HD motifs are globally linked to long-range regulation. (**A**) Position and evolutionary conservation of predicted TF motifs within the REX element. The conserved core of the REX element (chr16:51,624,707-51,624,984) is aligned with orthologous sequences from 12 mammalian species. Sequences matching TF binding preferences (below) are highlighted. (**B**) Schematic pipeline for genome-wide identification of putative enhancer-promoter interactions in the hindlimb. (**C**) Example of predicted E-P interaction between mHS919 limb enhancer and *Rad21*. The *Rad21* locus (chr15:5000000-5300000) is shown with Hi-C data (top) (Jiang et al., 2017), pseudobulk chromatin accessibility tracks and a violin plot for *Rad21* expression by cell type (bottom). Arches indicate hs919 enhancer-centric E–P interactions from enhancer capture Hi-C (red) or multiome (blue). **(D-E)** Table of top-most enriched limb-expressed TF motifs in long-range (400 kb – 2 Mb), as compared to short-range (10-200 kb) enhancers in developing limb buds for VISTA limb enhancers (**D**) and all predicted hindlimb enhancers (**E**) grouped by their similarity. [C/T]AATTA HD motifs are highlighted by red shading. False-discovery rate values are shown on the right. See **Table S2** for a complete list of motifs. (**F**) Distribution of enhancer-promoter distances for VISTA limb enhancers with 0-1 or ≥2 conserved [C/T]AATTA HD motifs (*P* = 0.0053, Wilcoxon rank sum test). Observation counts for each group are displayed in the outlined boxes at the base of the plot. (**G**) Representative long-range limb enhancers containing [C/T]AATTA HD motifs.

To investigate if LHX2, LHX9 and LEF1 motifs are also present next to other long-range limb enhancers, we used a collection of experimentally validated limb enhancers from the VISTA Enhancer Browser, a unique resource of human and mouse enhancers with *in vivo* activities characterized in transgenic mice (Pennacchio et al., 2006; Visel et al., 2007). We linked these *bona fide* limb enhancers to their putative target genes using the correlation between gene expression and open chromatin peaks from our single-cell ATAC-seq/RNA-seq experiment and E–P physical links from enhancer-capture-Hi-C experiments (Z. Chen et al., 2024) (**Fig. 5B, C**). We next performed motif analysis of these limb enhancers and included sequences adjacent to enhancers to account for potential neighboring REX elements (expanding the analyzed regions to at least 2 kb total length). We found eleven motifs that match binding preferences of limb-expressed TFs that are significantly differentially distributed between long-range acting (161 regions, 400 kb – 2 Mb E–P distance) and short-range acting (28 regions, 10 – 200 kb E–P distance) enhancer regions (FDR < 5×10^−2^ and Target Freq. > 30%) with LHX2 being the most significantly enriched (CAATTA, *P* < 3×10^−5^) (**Fig. 5D** and **Table S2**). HD TFs with consensus [C/T]AATTA binding preference comprised six of these eleven motifs. In contrast, short-range limb enhancers showed no enrichment for TF motifs over long-range enhancers.

We next examined all 19,276 limb mesenchyme E–P pairs predicted from our single-cell ATAC-seq/RNA-seq experiment (**Fig. 5E** and **Table S2**). Genome-wide, the LHX9 motif was the most significantly enriched in long-vs. short-range limb enhancers (CTAATTA, *P* < 3×10^−22^). Overall, [C/T]AATTA motifs, including both LHX2 and LHX9 motifs, comprised 15 out of the top 20 most significantly enriched motifs in long--range limb enhancers. The strong enrichment of [C/T]AATTA motifs in both sets of long-range limb enhancers suggests that their presence could be a defining feature of extreme-distance enhancers.

The REX element contains two conserved [C/T]AATTA motifs located within 30 bp from each other (**Fig. 5A**). To test if having more than one [C/T]AATTA motif could be characteristic of other long-range enhancers, we examined the number of [C/T]AATTA motifs in experimentally validated and multiome-predicted limb enhancers, extending the search space up to 2 kb to account for any potential REX elements (**Methods**). Limb enhancers containing more than one [C/T]AATTA motif within an extended 2 kb enhancer region were ∼500 kb farther (median distance) from their target promoters than other limb enhancers (*P* < 0.001 in validated; P < 0.05 in all; **Fig. 5F, G** and **S6B**). For example, the ZRS enhancer region contains four previously uncharacterized [C/T]AATTA motifs, three of which are evolutionary conserved across all jawed vertebrates (**Fig. 5G** and **6A**). Our results indicate that conserved [C/T]AATTA HD motifs are present in the REX element and are enriched in other extreme-distance limb enhancers genome-wide.

**Figure 6:**
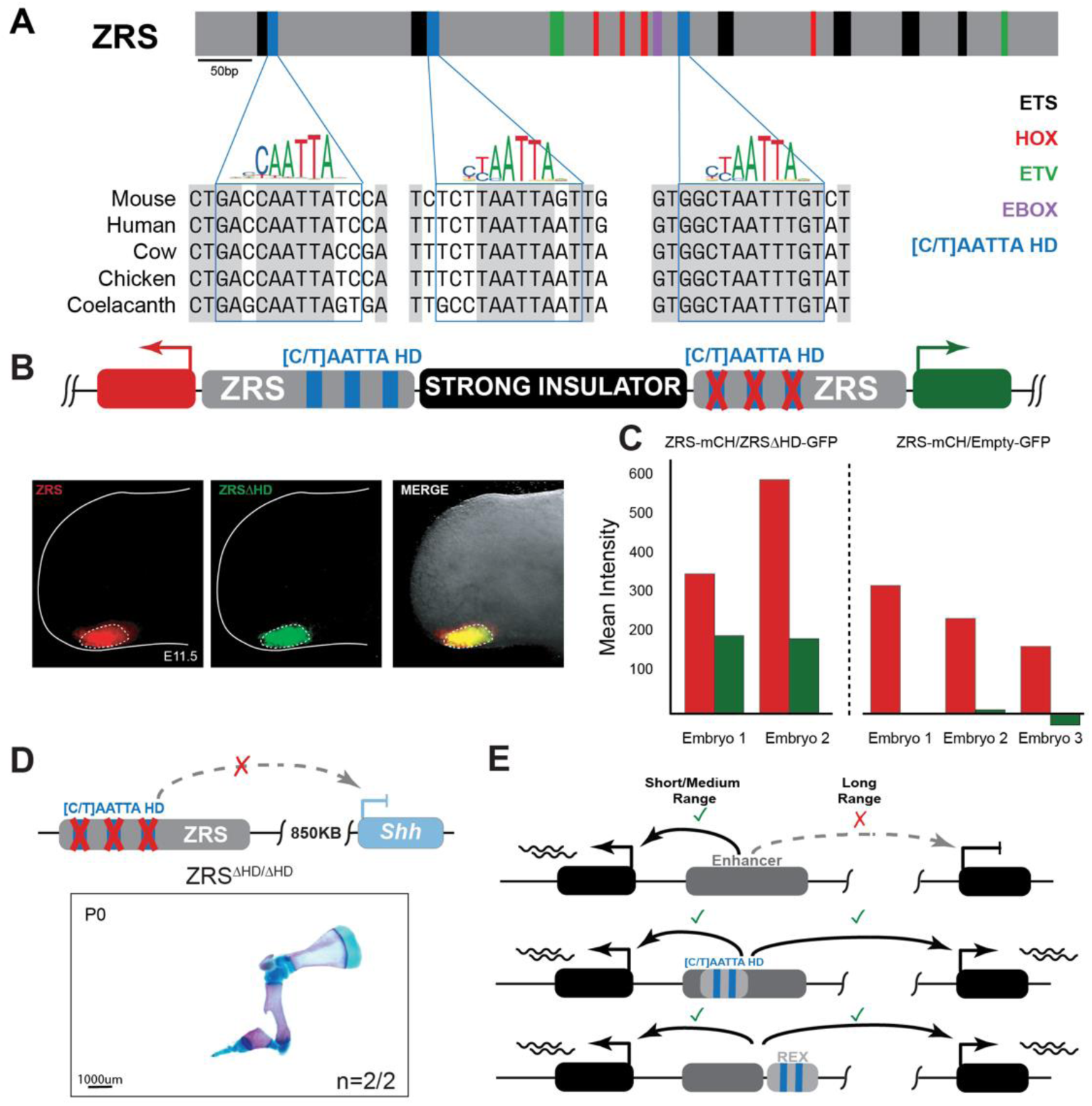
[C/T]AATTA HD motifs are required for long-range gene activation. (**A**) Position of TF binding sites previously identified (Kvon et al., 2016; Lettice et al., 2017, 2012) within the mouse ZRS core (chr5:29,314,881-29,315,667). Sequences matching [C/T]AATTA motifs binding preferences (below) are highlighted. Bump-outs show evolutionary conservation of [C/T]AATTA motifs. Blue boxes demarcate the mutagenized regions. (**B**) A dual-enSERT transgenic construct containing wild type mouse ZRS driving *mCherry* (red) and ZRS allele with mutated [C/T]AATTA HD motifs (ZRSΔHD) driving *eGFP* (green) separated by a strong synthetic insulator. Hindlimb bud images from a representative transgenic embryo are shown below. The white dotted line encircles the quantified region. (**C**) Quantification of normalized mean intensity in embryos containing ZRS-*mCherry*/ZRSΔHD-*eGFP* and control ZRS-*mCherry*/Empty-*eGFP* constructs. **(D)** Skeletal forelimb preparation from E18.5 mice with three mutated [C/T]AATTA sites within the endogenous ZRS enhancer (ZRSΔHD) (**E**) Proposed model of extreme long-range enhancer activation in the developing limb buds.

### [C/T]AATTA homeodomain motifs are required for long-range enhancer activity

To determine if [C/T]AATTA motifs are required for spatio-temporal limb-specific ZRS activity we employed dual-enSERT, our recently developed dual fluorescent reporter system which enables comparison of enhancer activities in live mice (Hollingsworth et al., 2023). We generated a construct that contains both a wild type ∼1.3 kb mouse ZRS allele upstream of *mCherry* and a mutant ∼1.3 kb mouse ZRS allele in which the three conserved [C/T]AATTA HD motifs are mutagenized (ZRSΔHD) upstream of *eGFP* separated by a strong insulator (SI). We left all other TF binding sites which were previously shown to be important for ZRS enhancer activity intact, including ETS1, ETV, HOX and E-BOX motifs (**Fig. 6A**). We injected the resulting ZRS-*mCherry*/SI/ZRSΔHD-*eGFP* bicistronic construct into mouse zygotes and collected transgenic embryos at E11.5. We detected mCherry and eGFP fluorescence in the ZPA, indicating that the ZRS enhancer lacking [C/T]AATTA motifs can act over short range and drive robust reporter expression (**Fig. 6B, C** and **S7C**). Importantly, we observed only mCherry expression and no detectable GFP expression in transgenic mice containing the ZRS-mCherry/SI/empty-eGFP construct, which do not have an enhancer driving *eGFP*, ruling out potential reporter crossactivation (**Fig. 6C** and **S7A, B**).

Having established that disruption of [C/T]AATTA motifs does not change limb enhancer activity in a reporter assay, we wondered if these motifs are specifically required for long-distance enhancer–promoter communication. We created a knock-in mouse line in which we disrupted the same three [C/T]AATTA motifs but now within the endogenous ZRS enhancer region, and again preserving all other known TF motifs. Strikingly, disrupting the [C/T]AATTA motifs alone resulted in a loss of limb outgrowth indistinguishable from complete loss of SHH function in the limb (**Fig. 1D** and **6D**). Taken together, our knock-in and transgenic results suggest that [C/T]AATTA HD motifs are critical for long-range ZRS activity but do not abolish its short-range activity.

## Conclusions

Here we have uncovered an evolutionarily conserved signature that is specifically required to confer long-distance enhancer–promoter communication. We also describe a prototypical element containing this signature, REX, that is sufficient to confer extreme-distance interactivity to heterologous short– and medium-range enhancers. This discovery arose from the observation that short– and medium-range developmental limb enhancers cannot initiate gene expression and support limb development when transplanted to a remote genomic location in place of a long-range enhancer. These results suggest that developmental enhancers with similar spatiotemporal activities are not always interchangeable at their remote endogenous genomic location and reveal a fundamental aspect of enhancer function that cannot be measured in traditional transgenic reporter experiments. Our results also provide a plausible mechanistic explanation for previous observations that linear distance can impact an enhancer’s ability to activate gene expression (Fukaya et al., 2016; Pachano et al., 2021; Rinzema et al., 2022; Swanson et al., 2010; Zuin et al., 2022).

Despite the failure to activate a remote gene, transplanted short– and medium-range enhancers can create an open chromatin structure similar to their endogenous location, indicating that the appropriate TFs are likely being recruited to the remote site. However, this chromatin environment is insufficient for long-range gene activation at the otherwise intact *Shh* locus. We identified REX as a novel modular *cis*-regulatory element adjacent to the HS72 limb enhancer at the *Sall1* locus and demonstrated that it is necessary and sufficient for long-range enhancer action. The REX element is not a classical enhancer, but its addition can extend the range of action of a heterologous short-range enhancer by more than 10-fold compared to its native range.

*Cis*-regulatory elements that facilitate enhancer activity continue to emerge as essential regulators of gene expression. This list includes *Drosophila* “tethering” (Calhoun & Levine, 2003; Calhoun et al., 2002; Levo et al., 2022) and “remote control” elements (Swanson et al., 2010) along with mammalian CTCF sites (L.-F. Chen et al., 2023; Despang et al., 2019; Kane et al., 2022; Lee et al., 2017; Paliou et al., 2019; Thiecke et al., 2020), CpG islands (Pachano et al., 2021) and enhancer booster elements (Blayney et al., 2023; Brosh et al., 2023; Hong et al., 2022). The REX element does not share sequence similarity with any of these elements, and it facilitates E–P activation over extreme distances, acting >840 kb from its target. Another key distinction is that rather than conferring robustness of remote enhancer activity, REX is both required and sufficient for long-range E–P activation and development of a key morphological structure.

Both the REX element and the ZRS contain highly conserved [C/T]AATTA binding sites. These sites match the binding preferences of homeodomain TFs, most notably LIM-homeodomain TFs LHX2 and LHX9. These HD motifs are also enriched in thousands of other long-range limb enhancers that act over distances greater than 400 kb, including the ZRS (**Fig. 5D-G**). Our experiments showing the preferential requirement of [C/T]AATTA motifs for long-but not short-range enhancer activity indicate that their cognate TFs could specifically mediate long-range E–P looping but might not be critical for general transcriptional activation. Indeed, LIM-domain-associated TFs such as GATA1, ISL1 and LHX2 are known to facilitate E–P looping at the β-globin locus as well as the assembly of interchromosomal olfactory gene compartments via their cofactor LIM-domain-binding factor (LDB1) (Deng et al., 2012; Krivega et al., 2014; Krivega & Dean, 2017; Lee et al., 2017; Liu & Dean, 2019; Monahan et al., 2019; Morcillo et al., 1997; Soler et al., 2010; Song et al., 2007). However, their general role in regulating long-range E–P interactions is poorly understood. Global enrichment of [C/T]AATTA HD motifs in extreme-distance enhancers suggests that there are likely more REX elements across the genome, with many of them integrated into the enhancer itself, as we showed with the ZRS (**Fig. 6E**). Further high-throughput investigation into long versus short-range enhancer activity will be necessary to understand how widespread REX elements are in the genome and across different cell types. Nevertheless, our results highlight that cell-type specificity and long-range activity are two distinct aspects of gene activation by enhancers, likely fulfilled by separate sets of TFs. This separation will have implications for interpretation of human genetics studies as some non-coding variants will likely affect only long-range enhancer ability while others will disrupt cell-type specific gene activation.

## Acknowledgments

The authors would like to acknowledge the UCI Transgenic Mouse Facility for help with the generation of the enhancer knock-in and transgenic mice, Dorota Skowronska-Krawczyk and Qianlan Xu for help with the ATAC-seq protocol. Funding: This work was supported by National Institutes of Health grants DP2GM149555 (to E.Z.K.), R01HG003988 (to L.A.P.), T32NS082174, T32GM008620 (supporting E.W.H.), and Ruth L. Kirschstein Predoctoral Individual NRSA fellowships F31HD112201 (to G.C.B.) and F30HD110233 (to E.W.H.). J.L-R. is supported by the Spanish MICINN through the PID2020-113497GB-I00 grant and the CEX2020-001088-M institutional grant. Research conducted at the E.O. Lawrence Berkeley National Laboratory was performed under Department of Energy Contract DE-AC02-05CH11231, University of California. This work was made possible, in part, through access to the Genomics High Throughput Facility Shared Resource of the Cancer Center Support Grant (CA-62203) at the University of California, Irvine and NIH shared instrumentation grants 1S10RR025496-01, 1S10OD010794-01, and 1S10OD021718-01.

## Author contributions

E.Z.K. conceived the project with input from G.B., D.E.D., A.V. and L.A.P. G.B. and E.Z.K. designed experiments. G.B. and E.Z.K. performed enhancer knock-in and transgenesis studies with help from S.J., B.C., K.C., M.L. and A.D. G.B. performed ATAC-seq experiments. A.F.B. analyzed the ATAC-seq data. Q.L. and D.S.K. helped develope ATAC-seq workflow. E.W.H. performed 10X multiome experiments and analyzed the data. E.W.H. and G.B. performed motif analysis. A.A.C. and J.L.R. performed ISH experiments. E.Z.K and G.B. wrote the manuscript with input from the remaining authors.

## Data and materials availability

All sequencing data is available at GEO: GSE243635.

## Methods

### Ethics statement

The animal work conducted in this study was reviewed and approved by the Lawrence Berkeley National Laboratory Animal Welfare and Research Committee and the University California Irvine Laboratory Animal Resources (ULAR) under protocols AUP-20-001 and AUP-23-005. Mice were housed in the animal facility, where their conditions were electronically monitored 24/7 with daily visual checks by technicians.

### Mouse tissue collections

The FVB/NCrl strain *Mus musculus* (Charles River) was used for all breeding experiments and mouse embryonic tissue collections. Mice were mated using a standard timed breeding strategy and E10.5, E11.5, E13.5 and E18.5 embryos along with P0 neonates were collected for staining or dissection using approved institutional protocols. Embryonic litters were kept on ice and processed one at a time to avoid degradation during collection. Embryos at unexpected developmental stages were excluded from the study.

### Generation of enhancer knock-in mice using CRISPR/Cas9

Knock-in mouse strains carrying replaced enhancer alleles were generated using a modified CRISPR/Cas9 pronuclear microinjection protocol (Z. Chen et al., 2024; Kvon et al., 2016). Briefly, we used previously described sgRNA and donor vectors containing heterologous enhancers surrounded by homology arms, both targeting the ZRS enhancer region (**Figs. S2A** and **S7D**) (Z. Chen et al., 2024; Kvon et al., 2016). Knock-in mice were generated by injecting a mix of Cas9 protein (IDT, final concentration of 25 ng/ul), sgRNA (IDT, 50 ng/ul) and donor plasmids (7.5-25 ng/ul) in an injection buffer (10 mM Tris, pH 7.5; 0.1 mM EDTA) into the cytoplasm of FVB embryos in accordance with standard procedures. Female mice (strain CD-1) were used as foster mothers. F0 mice were genotyped using PCR and Sanger sequencing (**Figs. S2B**) using primers F1, R1, F2 and R2 (see below and (Kvon et al., 2020)). Only founders carrying a single copy knock-in at the ZRS locus were used for breeding (Kvon et al., 2020). A knock-in mouse strain carrying part of the *lacZ* sequence in place of the ZRS was previously generated (Z. Chen et al., 2024).

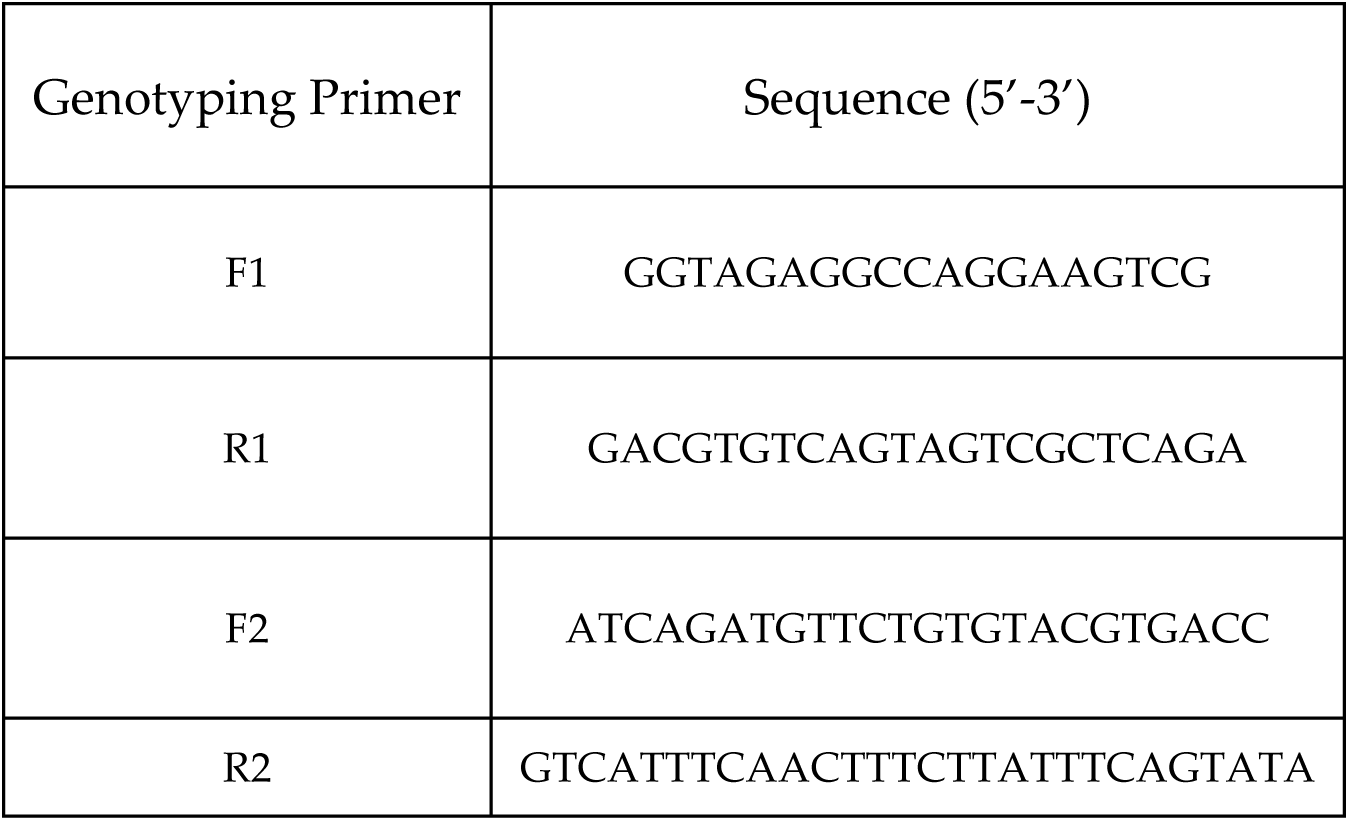

### Whole Mount *In Situ* Hybridization (WISH)

*Shh* transcript distribution in E10.5 and E11.5 mouse embryonic limb buds was assessed through whole mount in situ hybridization as previously described (Kvon et al., 2016). E10.5 or E11.5 Embryos were collected and fixed overnight in 4% paraformaldehyde, cleansed in PBT (PBS with 0.1% Tween-20) before being dehydrated through a methanol series to be preserved in 100% methanol. To perform ISH embryos were rehydrated, washed with PBT, and then bleached with 6% H_2_O_2_/PBT for 15 minutes before being washed again with PBT. Samples were then permeabilized for 15 minutes with 10 μg/ml of proteinase K in PBT, inactivated for 5 minutes in 2 mg/ml glycine/PBT, rinsed twice with PBT and re-fixed for 20 minutes 0.2% glutaraldehyde/4% PFA in PBT. Following fixation, embryos were with PBT (x3, 15mins) and incubated in prehybridization buffer (50% deionized formamide, 5× SSC, pH 4.5, 2% Roche Blocking Reagent, 0.1% Tween-20, 0.5% CHAPS, 50 mg ml−1 yeast RNA, 5 mM EDTA, 50 mg ml−1 heparin) for an hour at 70 °C, before overnight incubation in hybridization buffer containing 1 mg ml−1 Digoxigenin-labeled antisense riboprobes at 70 °C with gentle rotation. Dig-labeled riboprobes recognizing *Shh* were *in vitro* synthesized using RNA Labeling Mix (Roche) and T3 RNA polymerase (Roche). Next, embryos were washed multiple times in hybridization buffer with increasing concentrations of 2X SCC pH 4.5 finished with two washes with 2X SCC, 0.1% CHAPS. Embryos were next incubated for 45 minutes at 37C in 20 μg/ml RNase A in 2x SSC, 0.1% CHAPS before being washed in maleic acid buffer (100 mM Maleic acid disodium salt hydrate; 150mM NaCl; pH 7.5) (x2, 10min, RT and x2, 30min, 70C). Embryos were then transitioned to TBST (140mM NaCl; 2.7mM KCl; 25mM Tris-HCl; 1% Tween 20; pH 7.5) to prepare for an hour long blocking in 10% lamb serum/TBST and an overnight incubation at 4°C in a 1% lamb serum containing Anti-Dig-AP antibody (Roche, 1:5000). The next morning, embryos were washed with TBST (x3, 5min) to remove excess antibody and then further washed with TBST (x5, 1hr) before being equilibrated in NTMT (100mM NaCl, 100mM Tris-HCl; 50mM MgCl2; 1% Tween-20; pH 9.5). Alkaline phosphatase activity was detected by incubating in BM purple reagent (Roche) at room temperature in the dark with gentle agitation. The reaction was halted with PBT washes (x5, 10min) before transition to 4% paraformaldehyde in PBS for long-term storage. Embryos were imaged on a Flexacam C1 camera mounted on a Leica M125C stereomicroscope.

### Skeletal staining

Skeletal preparations were performed as previously described (Kvon et al., 2016). Embryos were harvested at E18.5 (or neonates were collected at P0) and decapitated before an overnight incubation in water at room temperature. Samples were then incubated in 65C water for 1 minute before the epidermis and organs were removed. Samples were fixed in 95% ethanol for storage before staining according to a standard Alcian blue/Alizarin red protocol (Rigueur & Lyons, 2014). Samples were incubated for at least 24 hours in Alcian blue stain (15% Alcian blue, 20% Acetic Acid in ethanol) followed by ethanol washes (x3) and an overnight ethanol incubation. Next, samples were cleared in 1% KOH for 20 minutes followed by counter-staining with Alizarin Red (5% Alizarin Red in 1% KOH) for 4 hours. Embryos were then further cleared with 1% KOH for 15 minutes followed by incubation in decreasing concentrations of 1% KOH and glycerol. The stained embryos were dissected in 80% glycerol and limbs were imaged at 1x using a ZEISS Stemi 508 microscope and a Axiocam 208 digital camera.

### LacZ Staining

LacZ staining was conducted as previously described (Kvon et al., 2020). Embryos were collected at E11.5 and fixed in 4% paraformaldehyde (PFA) for 30 minutes before washing (x3, 30 minutes) in embryo wash buffer (2mM MgCl2; 0.01% deoxycholate; 0.02% NP-40; 100mM phosphate buffer, pH 7.3). Embryos were then stained overnight in X-gal staining solution (0.8mg/ml X-gal; 4mM potassium ferrocyanide; 4mM potassium ferricyanide; 20mM Tris, pH 7.5 in wash buffer) to visualize LacZ activity. The next morning embryos were rinsed with PBS (x3, 10 minutes) and fixed again in 4% PFA. Images were recorded on a ZEISS Stemi 508 microscope with an Axiocam 208 digital camera.

### ATAC-seq library construction, sequencing and data analysis

Posterior limb bud tissue containing the ZPA region from E11.5 mouse embryos was dissected and manually dissociated, followed by snap freezing. After embryo genotyping, heterozygous embryos were processed for ATAC sequencing. Library construction was performed using the Omni-ATAC-seq protocol (Buenrostro et al., 2013, 2015), with an adjusted lysis step and transposition buffer based on the Omni-ATAC protocol (Corces et al., 2017). For the transposition reaction, nuclei were treated with Illumina Nextera transposase. The reaction was cleaned up using the Zymo DNA Clean and Concentrator kit, followed by preamplification of the transposed DNA using Nextera indexed primers with 5 cycles of PCR. The number of additional PCR cycles was determined by qPCR using 5ul of a partially amplified library. The resulting library was cleaned with AMPure XP beads (Beckman Coulter) and quantified by qPCR with Kapa Sybr Fast universal for Illumina Genome Analyzer kit. The library size was determined using the Bioanalyzer 2100 DNA High Sensitivity Chip (Agilent). The library was sequenced on the Illumina NovaSeq 6000 using 100 cycles and paired-end dual index read chemistry. The version of NovaSeq control software used was NVCS 1.6.0 with real-time analysis software RTA 3.4.4.

Fastq files were trimmed and clipped using trim_galore (https://www.bioinformatics.babraham.ac.uk/projects/trim_galore/) using the following options: –q 20 –-stringency 2 –-clip_R1 16 –-clip_R2 16. Reads were then aligned to the original mm10 genome assembly and a modified version of the genome containing the corresponding insertion using bedtools getfasta (Quinlan & Hall, 2010). Since the human HS72 enhancer inserted on chromosome 5 has an orthologous region on chromosome 8 of the mouse genome, we also mapped the reads only to chromosome 5 of the mm10 genome containing the insertion and only to chromosome 8 of the mm10 genome and could verify that no reads coming from the insertion were lost at the mouse orthologous region. Reads mapping to the positive strand were shifted by +4bp and reads mapping to the negative strand were shifted by –5bp to correct for the Tn5 insertion bias using the Unix command awk. Genomic tracks were generated using bedtools genomecov (Quinlan & Hall, 2010) by normalizing the mapped read counts to one million and dividing read counts by two except at the insertion sites.

### Single-cell multiomics and data analysis

Single-cell multiome ATAC-seq/RNA-seq was performed using a modified 10X protocol (10X Genomics, protocol CG000169 Rev D). Wild-type mouse embryos were harvested at embryonic day E11.5. A single hindlimb bud was dissected in ice-cold PBS and incubated with collagenase II (Gibco, #17101015, 0.2 uL at 100 u/ul) for 10 min at 37 C. Every five minutes, hindlimb tissue was triturated using a P200 pipette for mechanical dissociation into single cells. Immediately after collagenase treatment, 10% FBS (450 uL) was added and dissociated cells were spun down. The supernatant was removed, and cells were resuspended in 100 uL of ice-cold 10X nuclear lysis buffer (10X Genomics, protocol CG000169 Rev D) and incubated for 5 min on ice. Ice-cold wash buffer was added to the cells, lysed cells were centrifuged, and the supernatant was removed. Nuclei were resuspended in an ice-cold wash buffer, quantified, and inspected for viability using the Trypan Blue assay (Bio-Rad, 1450013). Nuclei were loaded at a concentration that would enable recovery of 10,000 nuclei by the 10X Chromium Single Cell Multiome ATAC + Gene Expression kit (10X Genomics, 1000285). Paired-end sequencing was performed on an Illumina NovaSeq 6000 for approximately 52,000 and 26,000 reads per cell for RNA– and ATAC-seq, respectively.

Fastq files were aligned to the mm10 genome assembly and barcodes were counted using CellRanger ARC (v2.0.2) (Satpathy et al., 2019). Expression count and chromatin peak matrices generated from CellRanger were further processed using the Signac R package (Stuart et al., 2021). The snATAC-seq dataset included a median of 31,000 reads and 14,200 high-quality fragments per cell, while the snRNA-seq dataset comprised a median of 68,000 unique molecular identifiers (UMIs) and 4,300 genes detected per cell. Low-quality nuclei were filtered out using standard Signac parameters and MACS2 was utilized for peak calling (Zhang et al., 2008). Transcriptome data were normalized and dimensionality was reduced using PCA (Seurat). DNA accessibility data were normalized using latent semantic indexing (LSI, Seurat). From the combined Seurat object, UMAP and nearest-neighbor analyses were performed to identify clusters with a resolution of 1 across twenty dimensions, as indicated by the ElbowPlot function, and based on their chromatin and gene expression profiles. Cell identities for the resulting sixteen clusters were assigned using well-known marker genes from the literature and past scRNA-seq datasets (Desanlis et al., 2020) (**Fig S1 and Table S2**).

### Enhancer–promoter assignment using multiome data

To link limb regulatory regions to their putative target genes, gene expression and open chromatin peaks (distance of 10 kb to 2 Mbp) were correlated using the LinkPeaks function from Signac (Stuart et al., 2021). This function accounts for bias in GC content (since Tn5 transposase is inherently biased toward cutting GC-rich regions), overall accessibility, and peak size. For analysis in **Figs. 1** and **5**, we used enhancers that were open (accessibility > 1) and genes that were expressed (normalized expression > 0.25) in mesenchymal cells.

### Motif Enrichment Analysis

A list of functionally-validated limb enhancers was obtained from the VISTA browser dataset (Visel et al., 2007). To assign target genes for these enhancers, we used the multiome E–P assignments or enhancer capture Hi-C data (Z. Chen et al., 2024). The resulting list was collapsed down into the longest distance interaction for each enhancer, and the pairs were separated into two groups based on the distance between the enhancer and target gene: short-range (10 – 200 kb) and long-range (400 kb – 2 Mb). To avoid excluding any potential REX elements, enhancers less than 2 kb were extended on each side to a distance of 2 kb. Only the longest-distance gene was considered for enhancers with multiple target genes to prevent statistical overrepresentation. Then, differential motif enrichment was then performed comparing short– and long-range E-P sets for (1) the bona fide E-P pairs (using VISTA enhancer coordinates) and (2) all predicted E-P pairs by multiome scATAC-seq/scRNA-seq. Differential motif analysis was performed using the findMotifsGenome.pl command in HOMER with a given size (Heinz et al., 2010) for long-range vs. short-range and vice versa. HOMER and JASPAR2022 (Castro-Mondragon et al., 2022) motifs were utilized in the motif search. Only motifs for limb-expressed TFs (curated from the hindlimb scRNA-seq (this study) and whole-limb bulk RNA-seq (Limb-Enhancer Genie (Monti et al., 2017)) were considered in the analysis.

To analyze enrichment of [C/T]AATTA HD motifs within long-range vs. short-range enhancers we used the findMotifsGenome.pl command from HOMER with the given size in order to generate motif occurrences using MA0700.2.LHX2 and LHX9(Homeobox)

/Hct116-LHX9.V5-ChIP-Seq(GSE116822) PWMs. The Genomic Scores package (Puigdevall & Castelo, 2018) was using in R with the phastCons60way.UCSC.mm10 dataset in order to filter motif occurrences to a conservation score of greater than zero, and then overlapping motif occurrences were collapsed into one region for subsequent analysis. Statistical significance was calculated using a wilcoxon test.

To localize [C/T]AATTA HD motifs within individual enhancers, the findMotifsGenome.pl command in HOMER was used with the region size set to the average size of the enhancers tested. The output motifs were then mapped back to genomic coordinates. The conservation of each motif occurrence was analyzed using the Table Browser (Karolchik et al., 2004) tool with 100 Vert. Cons (phyloP100wayAll) used for human sequences and Vertebrate Cons (phyloP30wayAll) used for mouse sequences.

### [C/T]AATTA HD motif mutagenesis

For [C/T]AATTA motif mutagenesis, the [C/T]AATTA HD motif regions within the ZRS were replaced with random sequences. FIMO (Grant et al., 2011) was used to ensure that the mutagenized sequence does not have any [C/T]AATTA motif matches. Changes were as follows: GGATAATTGGTC > CGACGTCTGTAG (1st HD site), AACTAATTAAGA > TCGACGGACACT (2nd HD site), and GACAAATTAGCC > AGGGACTGCTCT (3rd HD site). The resulting ∼1.3 kb ZRSΔHD sequence was used for dual-enSERT and knock-in analysis (**Figs. 6** and **S7**).

Presence of motifs in knock-in mice was confirmed through PCR using primers F1, R1, F2, R2, F3 and R3 (below and **Figs. S2A** and **S7D-E**) and sanger sequencing.

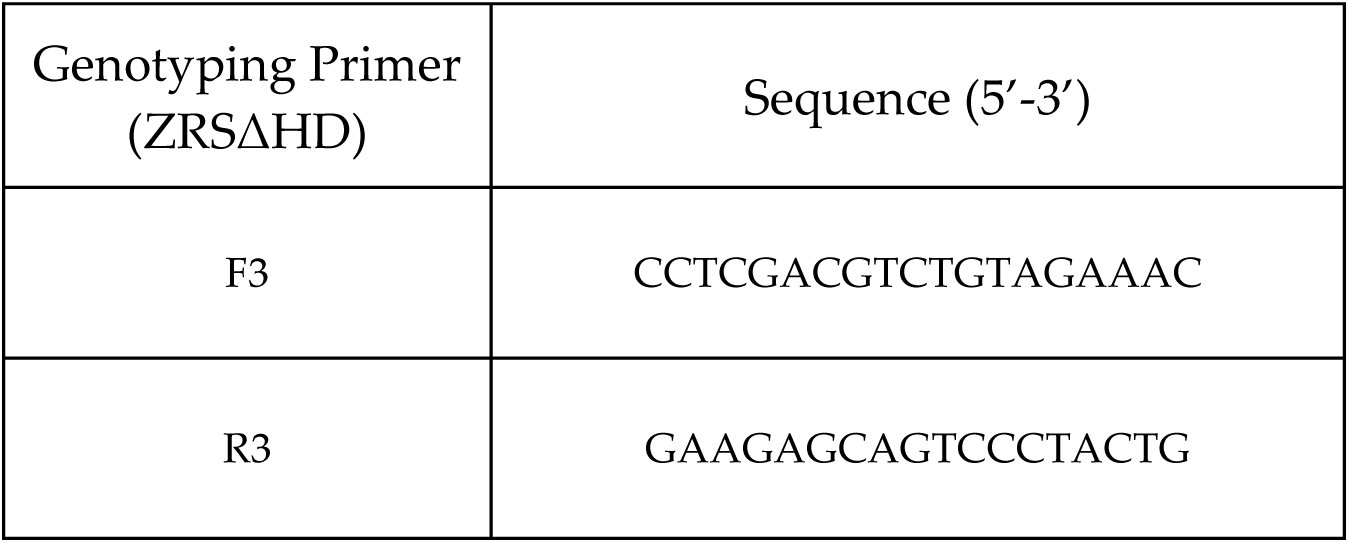

### Live fluorescent imaging and quantification

Imaging of dual-enSERT embryos was conducted as previously described (Hollingsworth et al., 2023). Embryos were imaged using a Zeiss V20 stereoscope equipped with a monochromatic camera (Axiocam 202, Zeiss), fiber optic light source (Zeiss, CL1500), and LED fluorescent laser (X-Cite, Xylis) emitting at 488 (eGFP) and 555 (mCherry) nm wavelengths. Merged images were created using the Zeiss Biolite Software.

For quantification, .czi files were imported in Fiji (Schindelin et al., 2012). The region of interest, the ZPA, was defined around the brightest region of mCherry signal and applied to the GFP channel. A control region was anterior limb bud was also delineated to determine the background fluorescence of the embryo in each channel. Mean intensity was measured within both regions for both channels. The measurement for the background was subtracted from the region of interest in order to generate the mean intensity.

### Generation of knock-in and transgenic reporter mice

All constructs in this study were generated using Gibson Assembly cloning (New England Biolabs). The targeting vector containing heterologous enhancers (e.g., ZRS-MHA-PCR4-HS72 for HS72 enhance) was created by Gibson cloning of the heterologous enhancers into the previously described ZRS targeting vector (Kvon et al., 2020).

The HS72-*Shh*-promoter::*lacZ* construct was created by introducing the HS72 enhancer (chr16: 51,623,899 – 51,624,805; hg38) into the *hsp68*-promoter-*lacZ* reporter vector (Kvon et al., 2016) followed by replacement of the *hsp68* promoter with the mouse *Shh* promoter (chr5: 28,466,764 – 28,467,284; mm10). The REX-*Shh*-promoter::*lacZ* construct was created by introducing the REX element into the PCR4-*Shh*-promoter::*lacZ*-H11 vector (Addgene Plasmid #139098) (Kvon et al., 2020). The sequence used for the REX element was expanded slightly to include the entire conserved block (**Fig S5A**) in order to more thoroughly test the enhancer activity of the fragment. The HS72-*Shh*-promoter::*Shh* construct was created by cloning the HS72 enhancer followed by the mouse *Shh* promoter and *Shh* ORF into the PCR4-*Shh*-promoter::*lacZ*-H11 vector in the opposite orientation relative to the *lacZ* reporter gene. ZRS_mCH/ZRSΔHD_eGFP-SuperIns was generated by inserting wild type ZRS and ZRSΔHD sequences into the *dual-enSERT-2.2* plasmid (Hollingsworth et al., 2023).

Transgenic mice carrying enhancer-reporter transgenes were created using random transgenesis (HS72-*Shh*-promoter::*lacZ* construct) or site-directed enSERT transgenesis (REX-*Shh*-promoter::*lacZ,* HS72-*Shh*-promoter::*Shh*, and ZRS_mCH/ZRSΔHD_eGFP-SuperIns constructs) as previously described (Kothary et al., 1989; Kvon et al., 2020; Pennacchio et al., 2006). After pronuclear microinjections (below), F0 embryos were harvested at embryonic day E11.5 and processed for LacZ staining (HS72-*Shh*-promoter::*lacZ* and REX-*Shh*-promoter::*lacZ*) or fluorescent imaging (ZRS_mCH/ZRSΔHD_eGFP-SuperIns) or at embryonic day E13.5 and analyzed for limb morphology (HS72-*Shh*-promoter::*Shh*). The embryos were genotyped by PCR and Sanger sequencing as previously described (Kvon et al., 2020). The sequences are detailed below (**Fig. S8**)

## 1. ZRS-MHA-PCR4-HS72 (Targeting Vector)

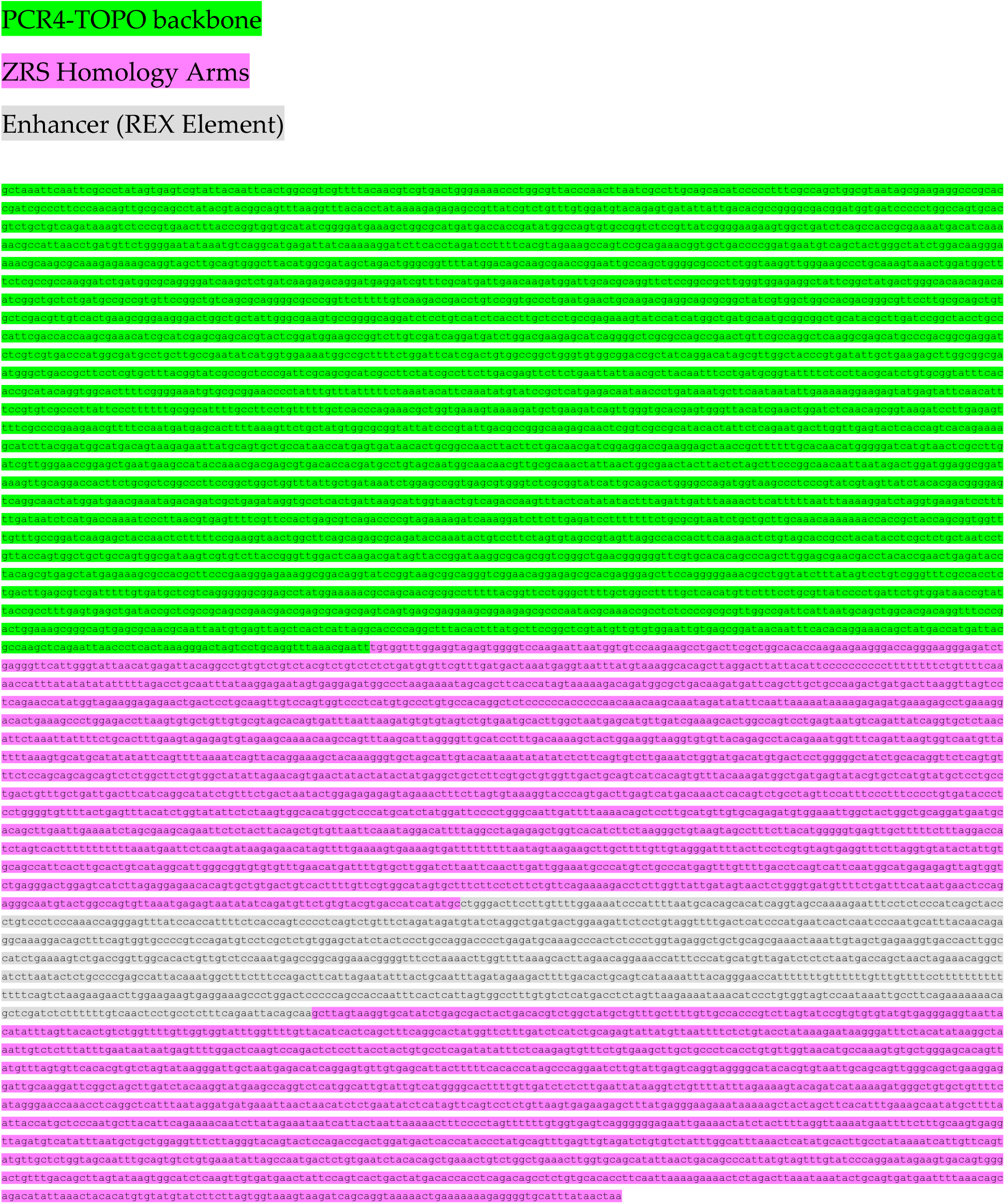

## 2. REX-*Shh*-promoter::*lacZ*

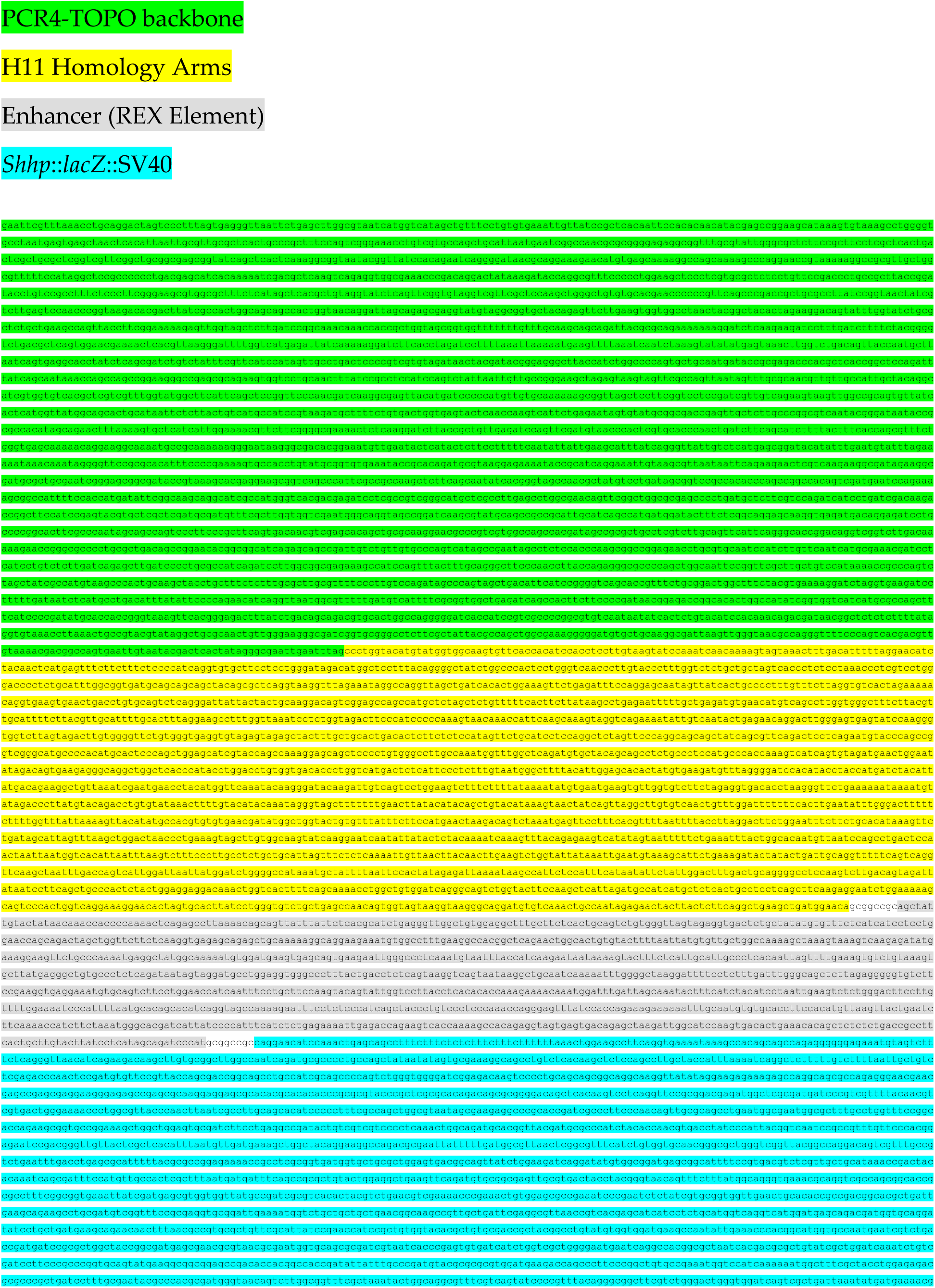
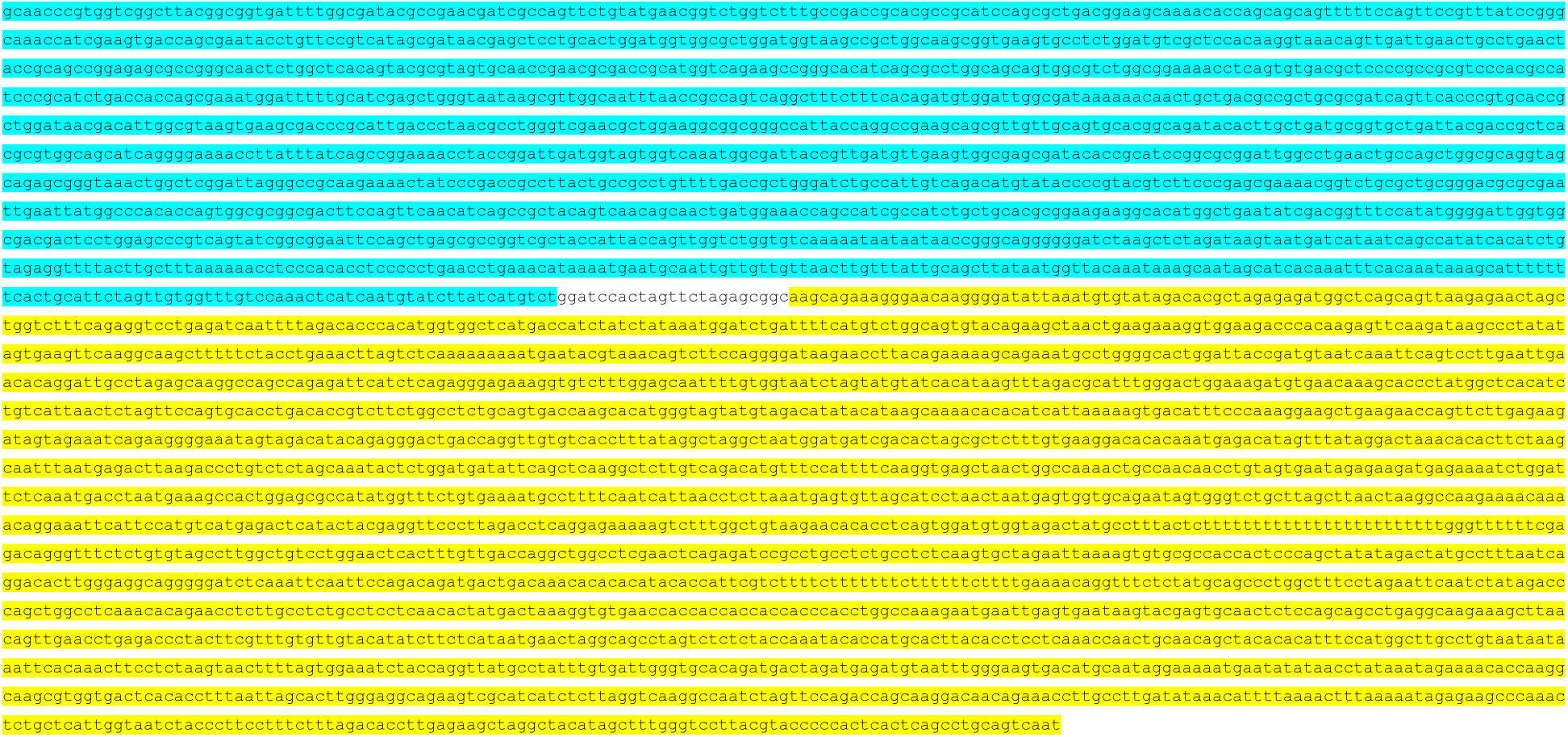

## 3. HS72-*Shh*-promoter::*lacZ*

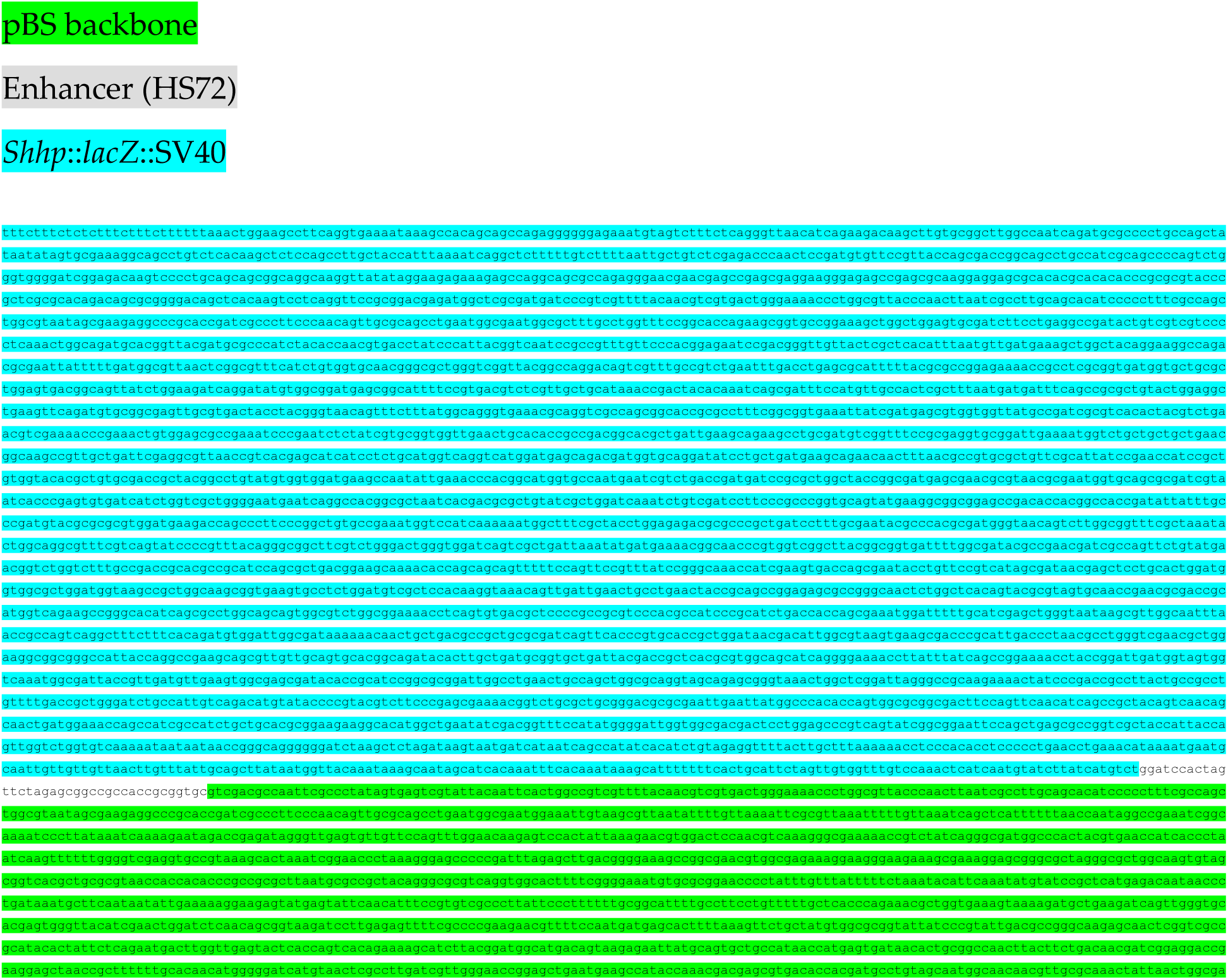
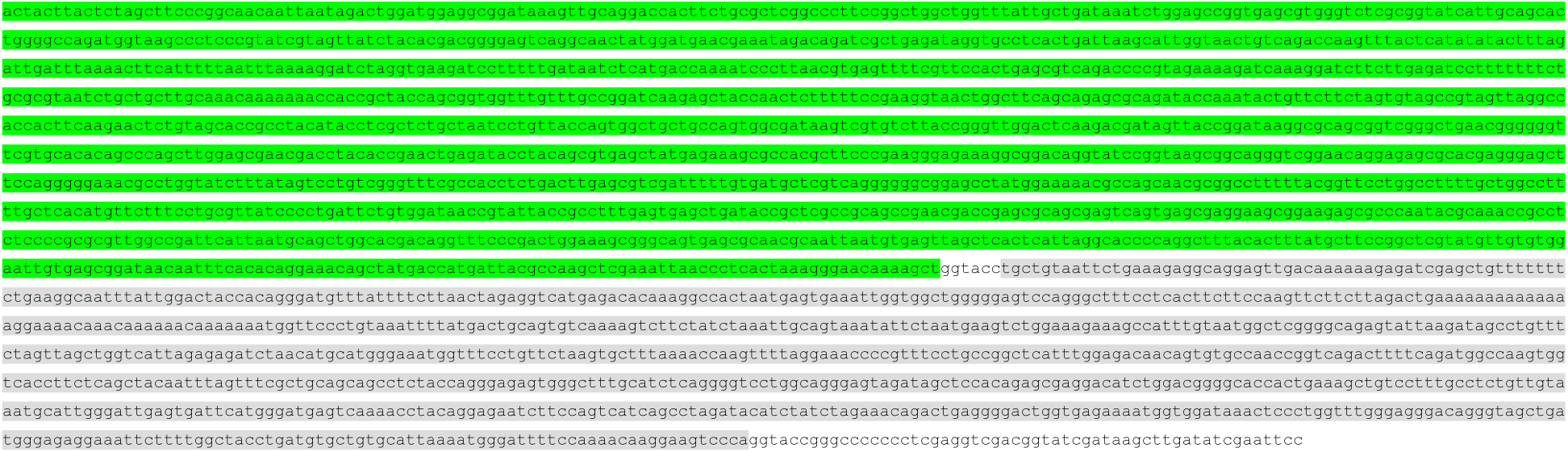

## 4. HS72-*Shh*-promoter::*Shh*

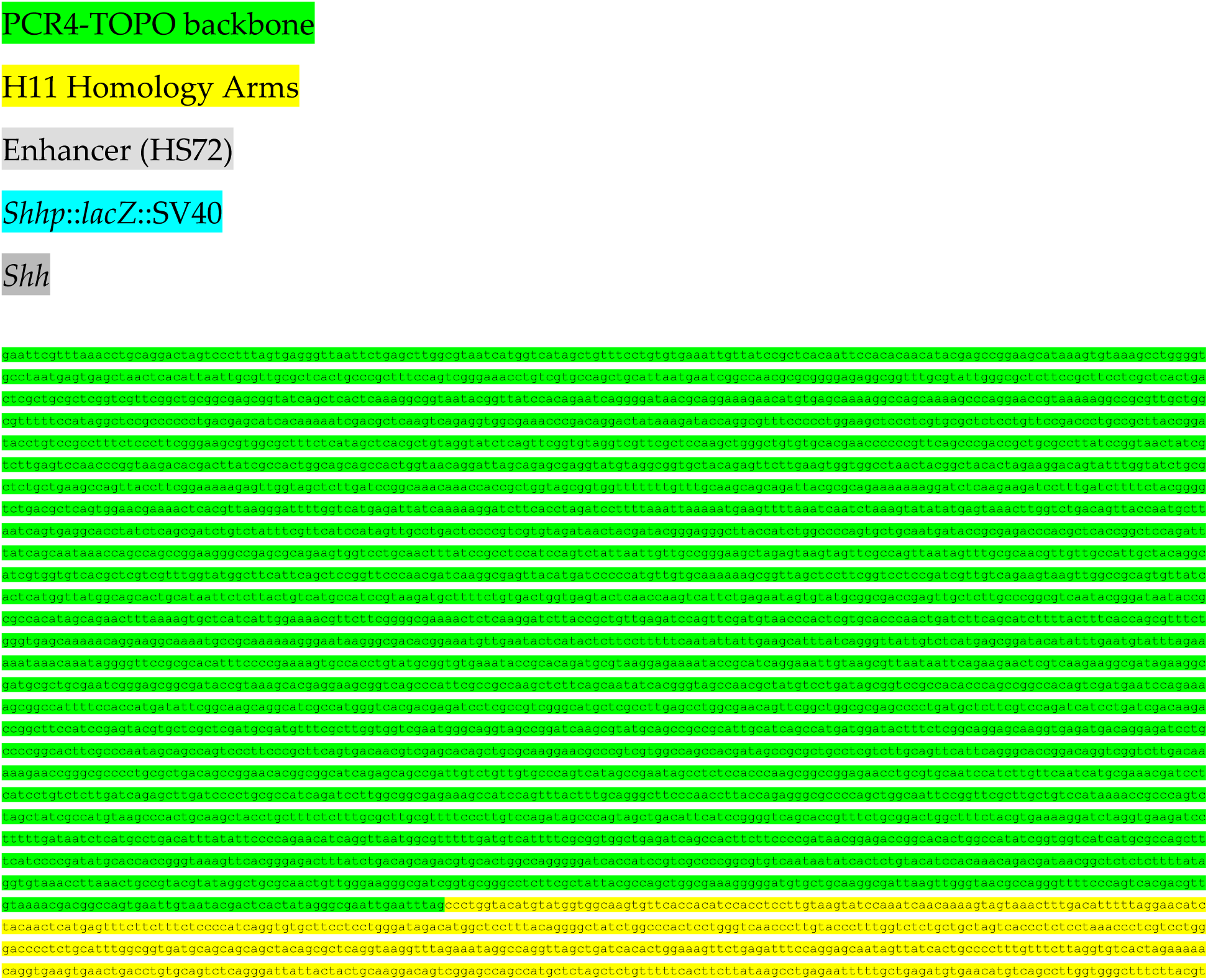
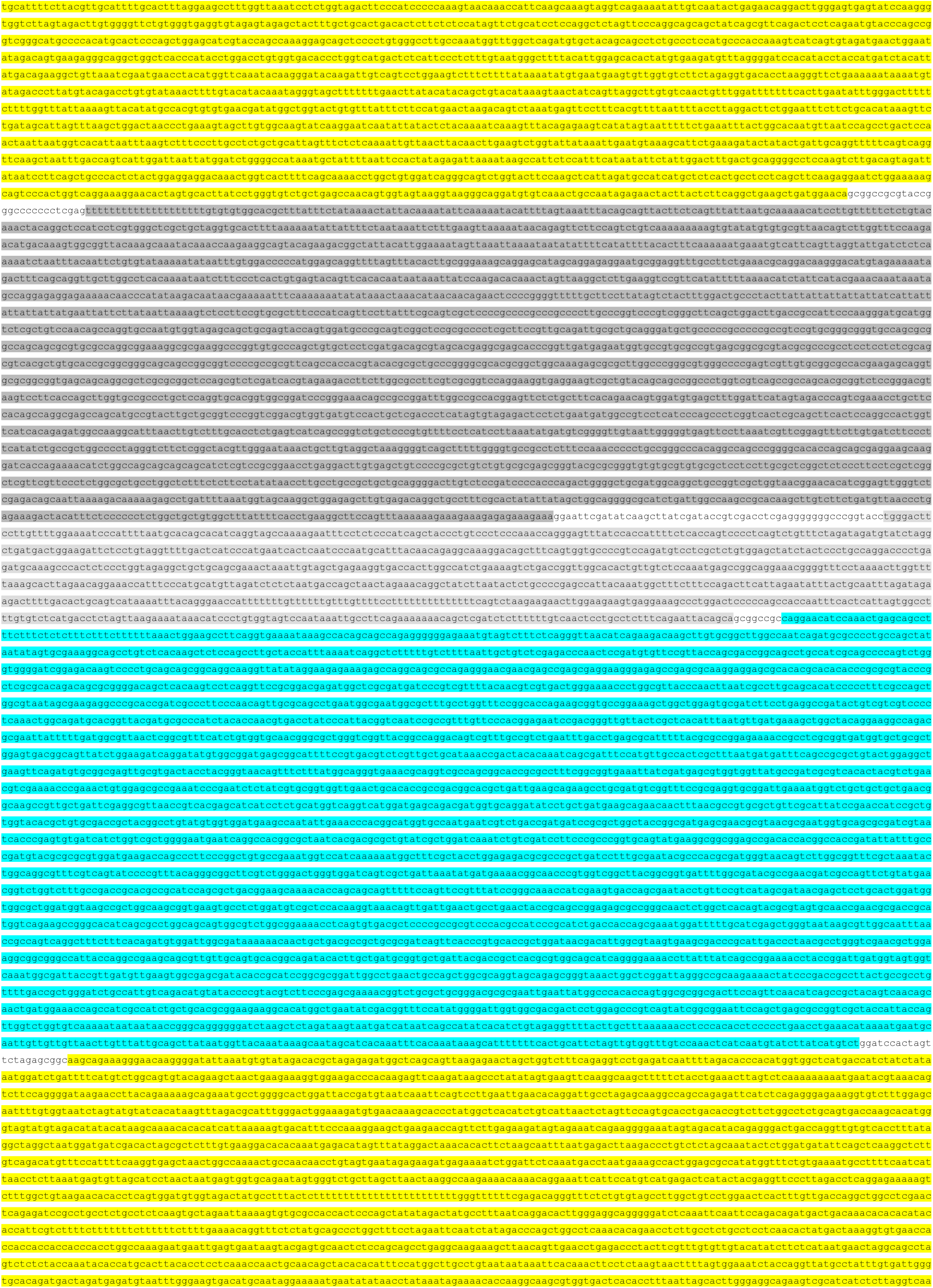
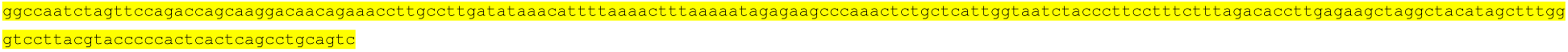

## 4. ZRS_mCH/ZRSΔHD_eGFP-SuperIns

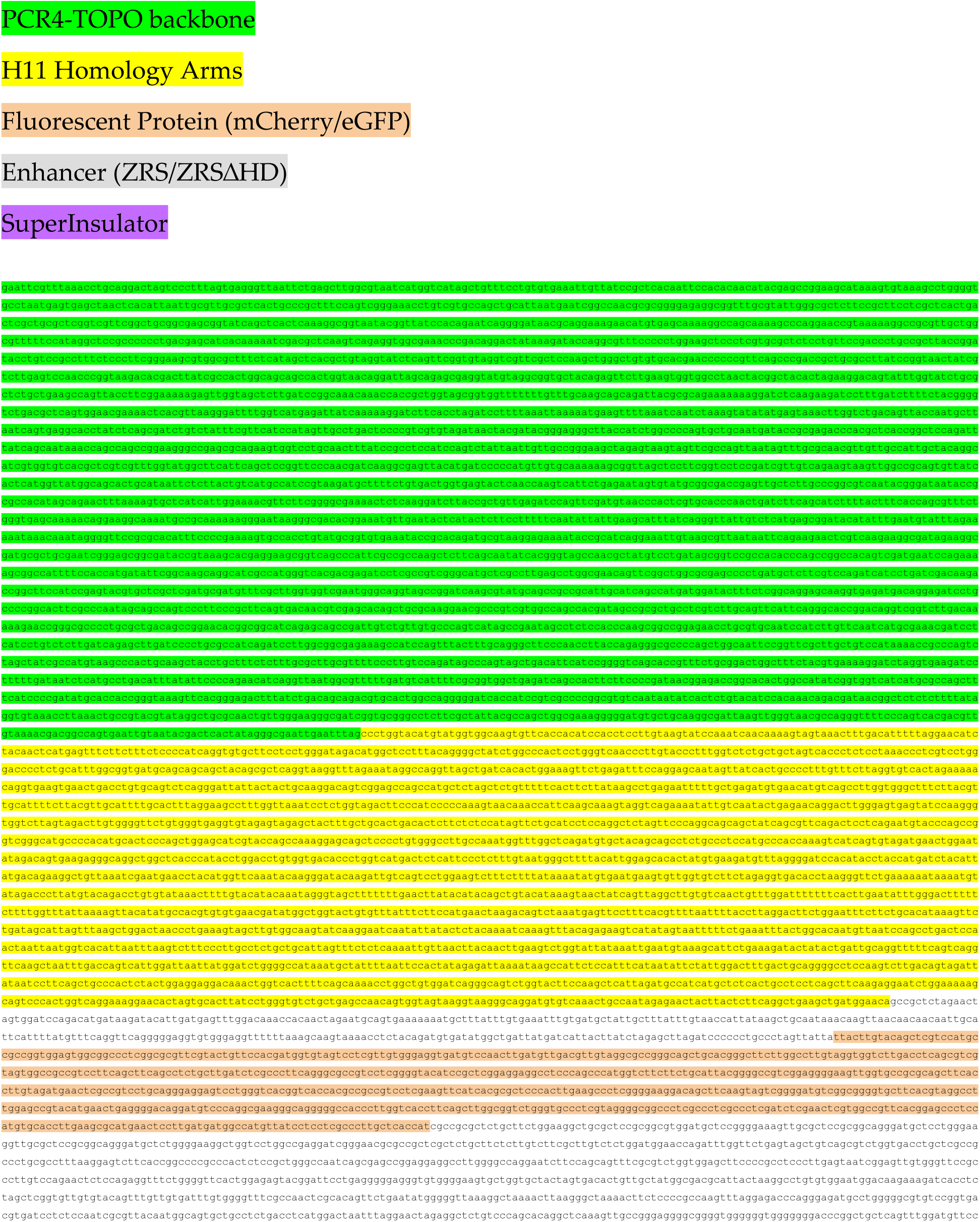
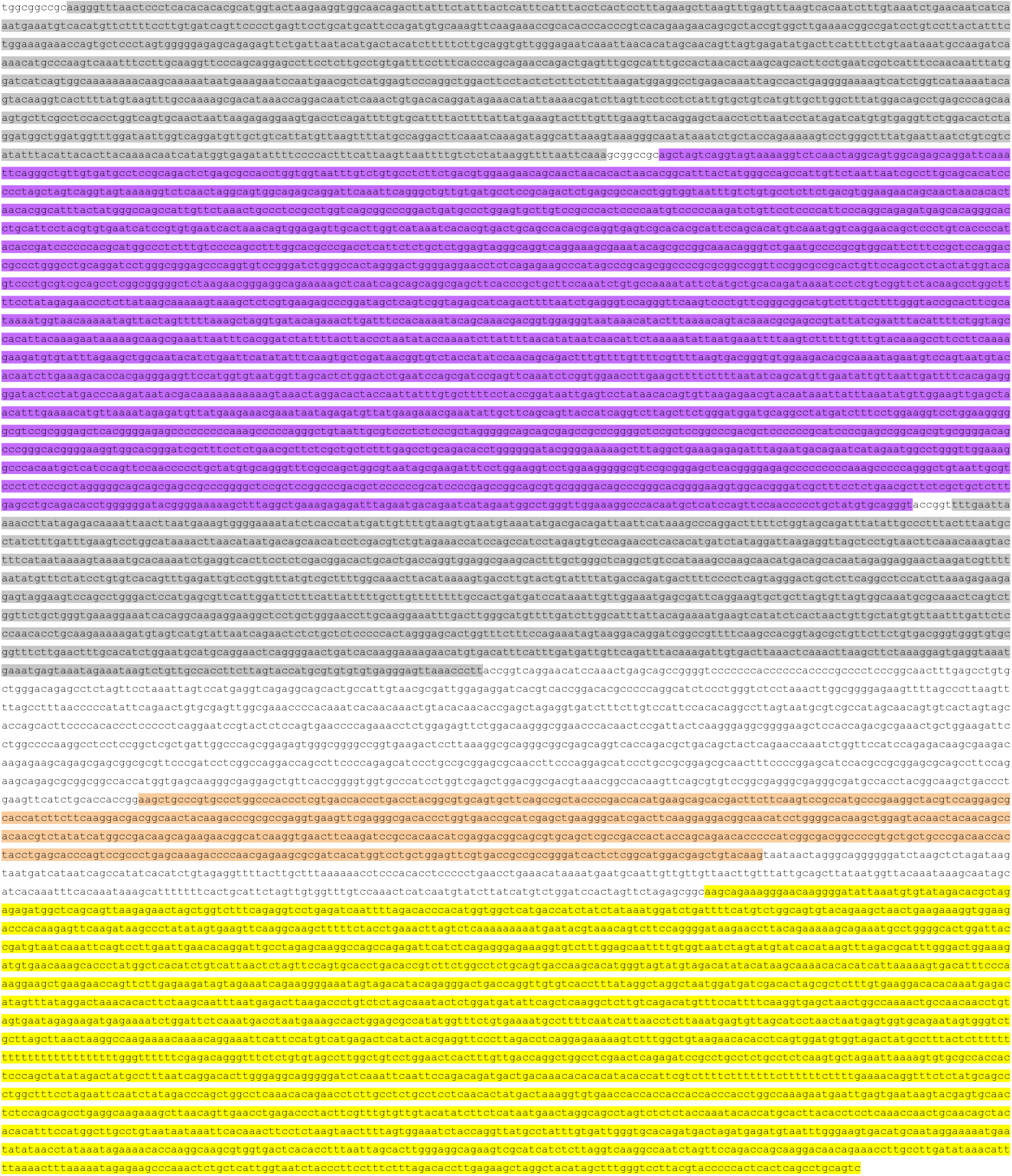

### Statistics and reproducibility

No prior analyses were used to determine sample size before experimentation. Embryos that were not at the correct developmental stage were excluded from data collection. For LacZ and skeletal Alcian blue/Alizarian red staining, embryos were identified by number and researchers were blinded to genotype. To determine the statistical significance of [C/T]AATTA motif enrichment, Wilcoxon rank-sum tests were used.

## Supplementary Figures

**Supplementary Figure 1:**
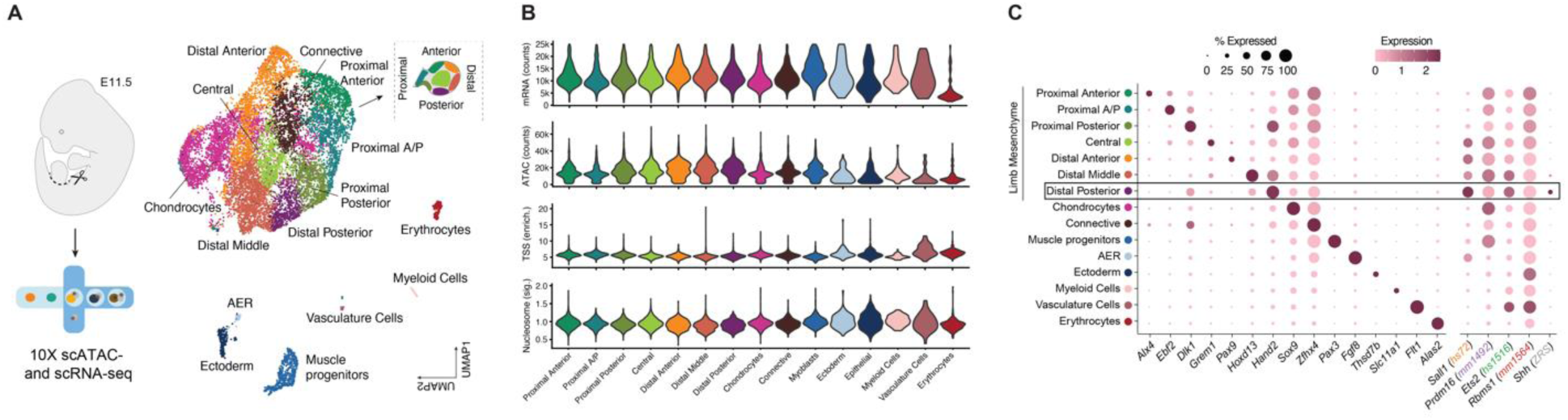
Multiome analysis (scATAC-seq + scRNAseq) in a mouse hindlimb bud. (**A**) Single-cell multiomics in developing E11.5 mouse hindlimb bud. The UMAP plot depicts cell type clusters based on integrated scRNA-seq and scATAC-seq data, along with a cartoon mapping the different regions of the hindlimb bud. (**B**) Single-cell multi-omics quality control violin plots for a number of mRNA counts, ATAC read counts, transcription start site (TSS) enrichment, and nucleosome signal by cell type cluster. (**C**) Dot plot showing expression of cell-type-cluster-specific genes (left) and putative target genes of enhancers tested in this study (right) across cell clusters. Color represents expression level and the size of the circle depicts the percent of cells expressing each gene. The boxed region (Distal Posterior) encompasses the posterior limb bud region containing the Zone of Polarizing Activity (ZPA), where *the ZRS normally activates Shh*. Apical ectodermal ridge (AER).

**Supplementary Figure 2:**
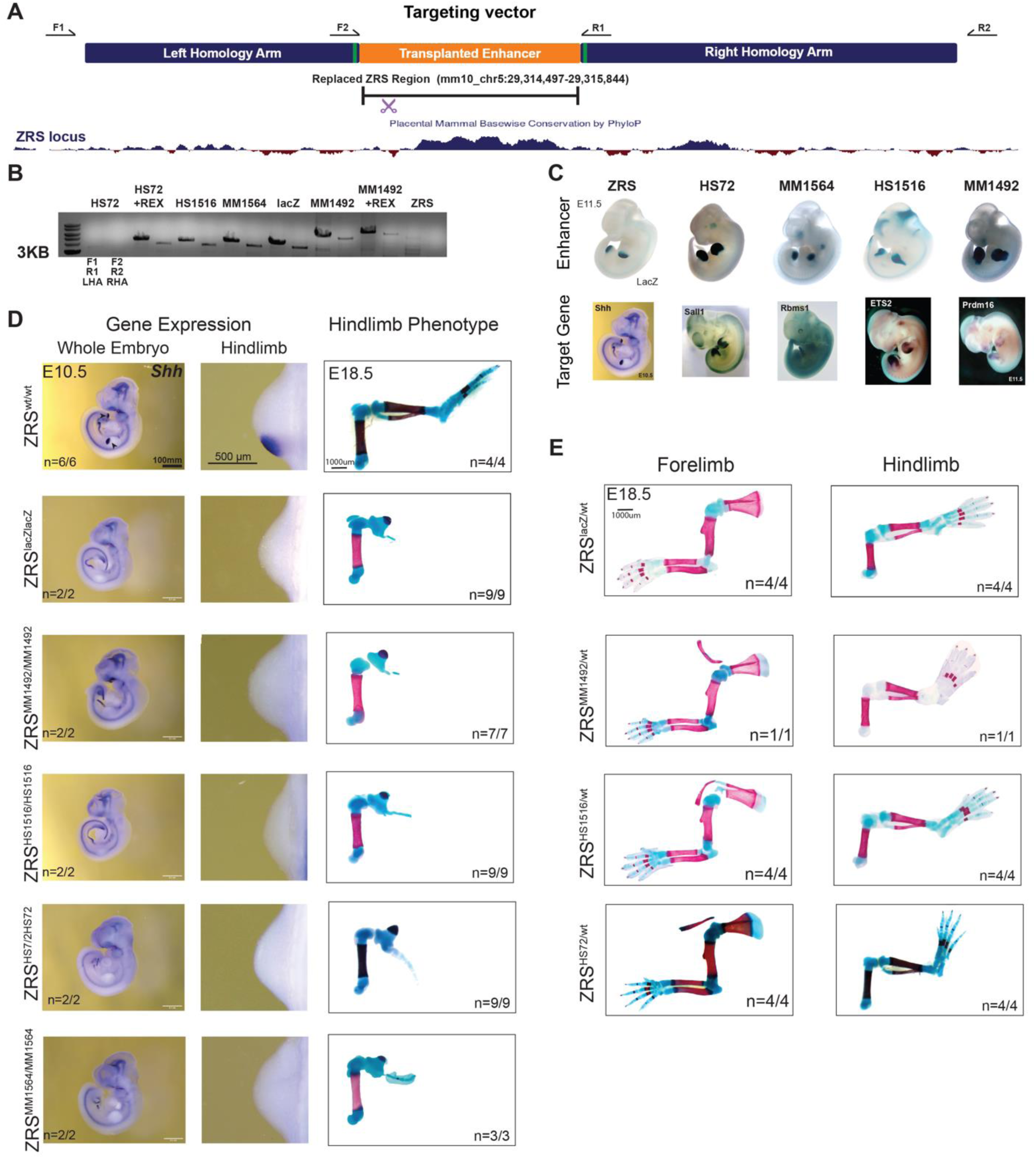
Generation and characterization of mice with transplanted enhancers. (**A**) Schematic overview of enhancer replacement strategy. A 4.5 kb mouse genomic region containing the ZRS enhancer is shown together with the vertebrate phyloP conservation (dark blue). The donor vector contained two homology arms with vector-specific sequences for genotyping (green) and a corresponding replaced region containing the transplanted enhancer and mutagenized sgRNA recognition site (purple, 5’-agtaccatgcgtgtgtTtTagCC-3’). PCR primers used for genotyping are shown as arrows. See **Methods** and Kvon et al. 2020 for more details. (**B**) Shown are the results of PCR genotyping for each knock-in mouse line. (**C**) Top: E11.5 enhancer activity for ZRS, HS72, MM1564, HS1516 and MM1492 enhancer. Bottom: whole mount *in situ* hybridization for *Prmd16* (Shimizu et al., 2013) (MM1492 target gene), *ETS2* (Shimizu et al., 2013) (HS1516), *Rbms1* (Groza et al., 2023) (hs1564), *Sall1* (Nishinakamura et al., 2001) (HS72), and *Shh* (ZRS) in E10.5-E11.5 mouse embryos. (**D**) *Shh* mRNA whole mount *in situ* hybridization analysis in wild type and knock-in mouse embryos (first two columns). The corresponding hind limb skeletal preparations of E18.5 wild type and knock-in mouse embryos are shown (third column). The number of embryos that exhibited representative limb phenotype over the total number of embryos with the genotype is indicated. Genotypes for each row are displayed on the left. (**E**) E18.5 skeletal staining showing the forelimb (left panel) and hindlimb (right panel) phenotype in heterozygous knock-in embryos.

**Supplementary Figure 3:**
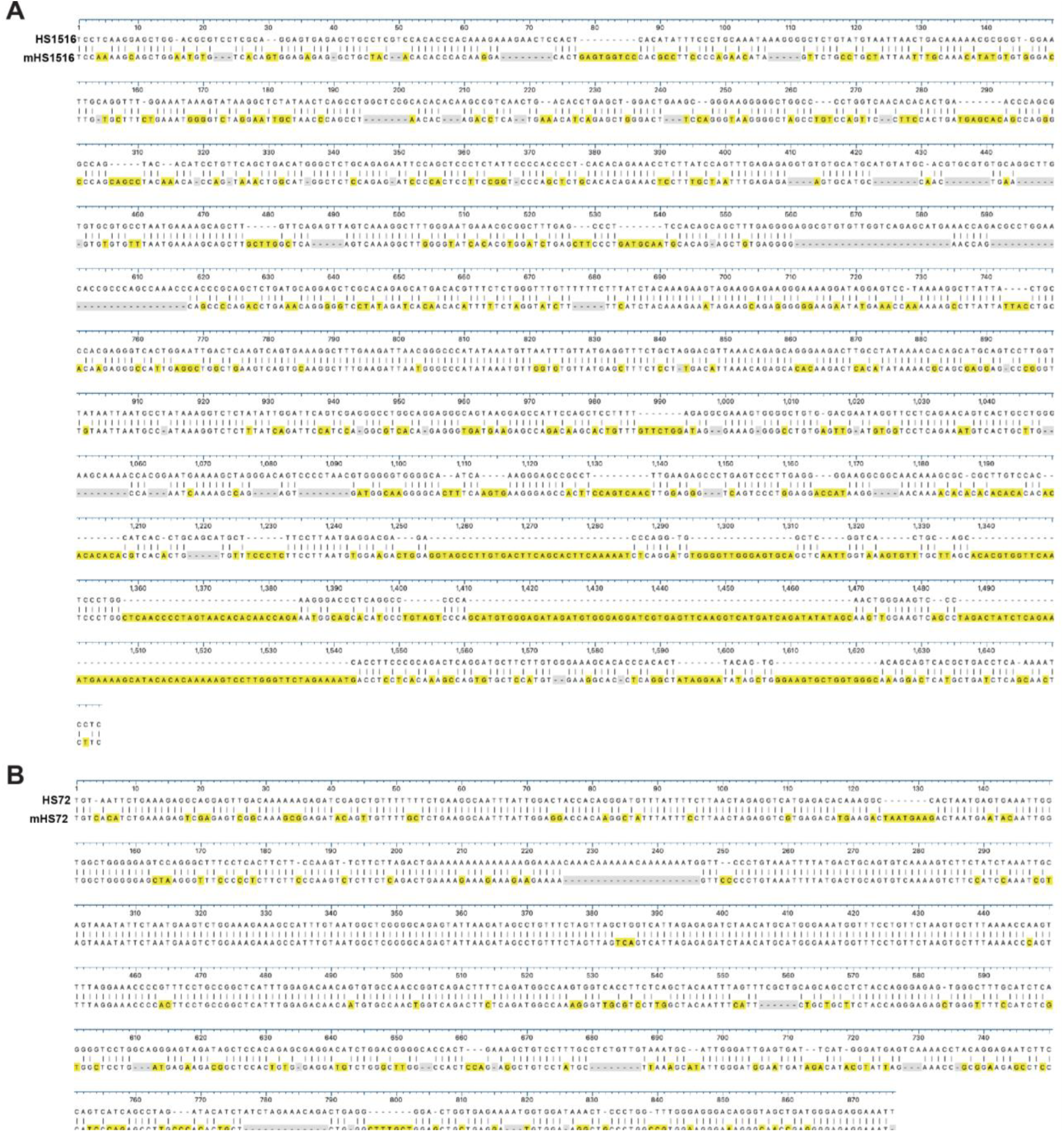
Pairwise alignment of the human HS72 and HS1516 enhancer sequences with their respective mouse homologues.

**Supplementary Figure 4:**
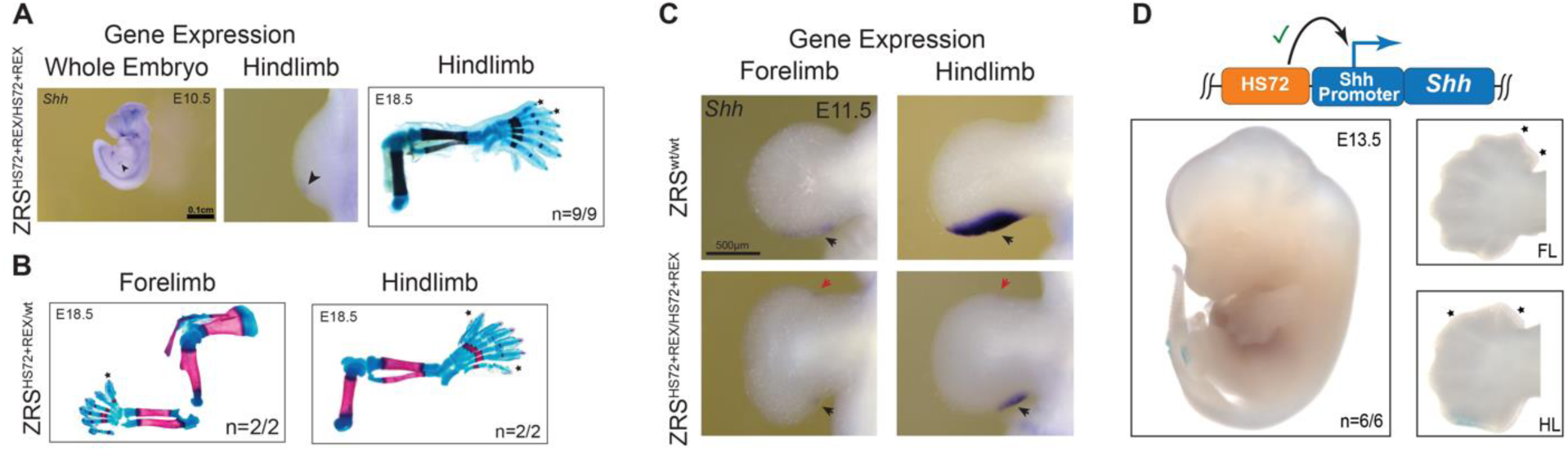
The REX element is required for long-range HS72 enhancer activity in the knock-in mice. (**A**) Gene expression analysis of *Shh* using mRNA whole mount *in situ* hybridization analysis in homozygous ZRS^HS72REX/HS72REX^ knock-in mouse embryos at E10.5 (left two panels). Arrows point to the *Shh* expression in the ZPA. The corresponding hindlimb skeletal preparations at E18.5 are shown (third column). The number of embryos that exhibited representative limb phenotype over the total number of embryos with the genotype is indicated. (**B**) Forelimb (first panel) and hindlimb (second panel) skeletal phenotypes in heterozygous ZRS^HS72REX/+^ knock-in embryos at E18.5. (**C**) Comparative *Shh* mRNA *in situ* hybridization analysis in wild type (top row) and homozygous ZRS^HS72REX/HS72REX^ knock-in (bottom row) mouse embryos during limb bud development in E11.5 embryos. Black arrows point to areas of *Shh* expression in the ZPA. Red arrows point to ectopic expression in the anterior portion of the limb bud. * Extra digits. (**D**) Transgenic E13.5 mouse embryo with the HS72 limb enhancer placed upstream of the *Shh* promoter and ORF and integrated at H11 safe-harbor locus (light blue). Close-up images of limbs are shown on the right. Asterisks indicate extra digits (polydactyly). Numbers of embryos with limb polydactyly in both forelimb and hindlimb buds over the total number of transgenic embryos screened are shown. FL, forelimb. HL, hindlimb.

**Supplementary Figure 5:**
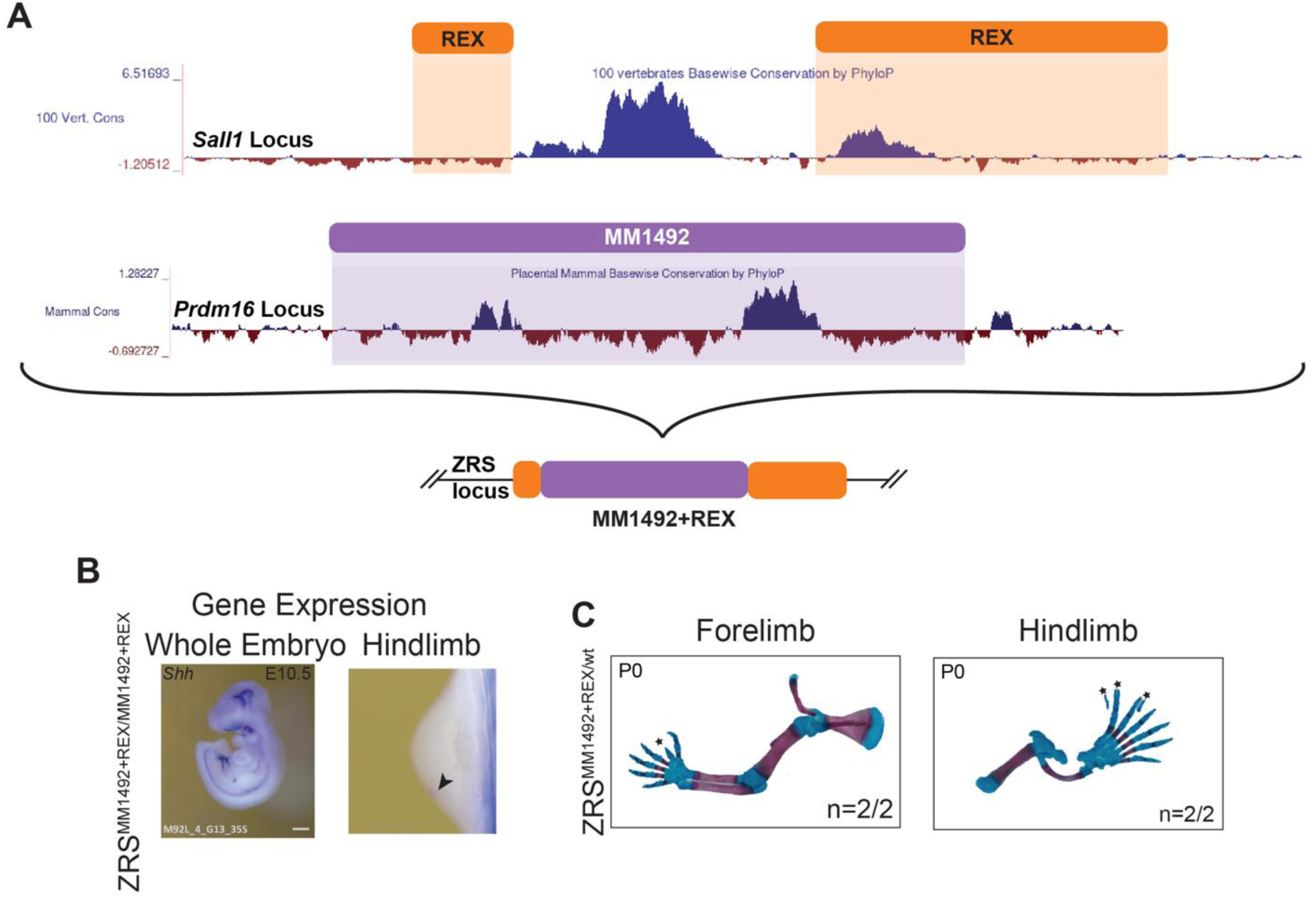
The addition of the REX element to the MM1492 limb enhancer is sufficient to rescue distal enhancer activity and *Shh* expression. (**A**) Schematic showing the sequences from the HS72 (chr16:51623658-51623900 and chr16:51624689-51625572; hg38) and MM1492 (chr4:154706480-154712089; mm10) loci that make up the chimeric MM1492+REX element. The orange bars indicate the expanded region flanking the core HS72 enhancer that contains the REX element. The purple bar marks the MM1492 enhancer sequence. The core HS72 enhancer was swapped out for the MM1492 enhancer to generate the MM1492+REX construct (bottom). (**B**) Gene expression analysis of *Shh* using mRNA whole mount *in situ* hybridization analysis in homozygous ZRS^MM1492REX/MM1492REX^ knock-in mouse embryos at E10.5. (**C**) Forelimb and hindlimb skeletal phenotypes of heterozygous ZRS^MM1492REX/+^ embryos at E18.5.

**Supplementary Figure 6:**
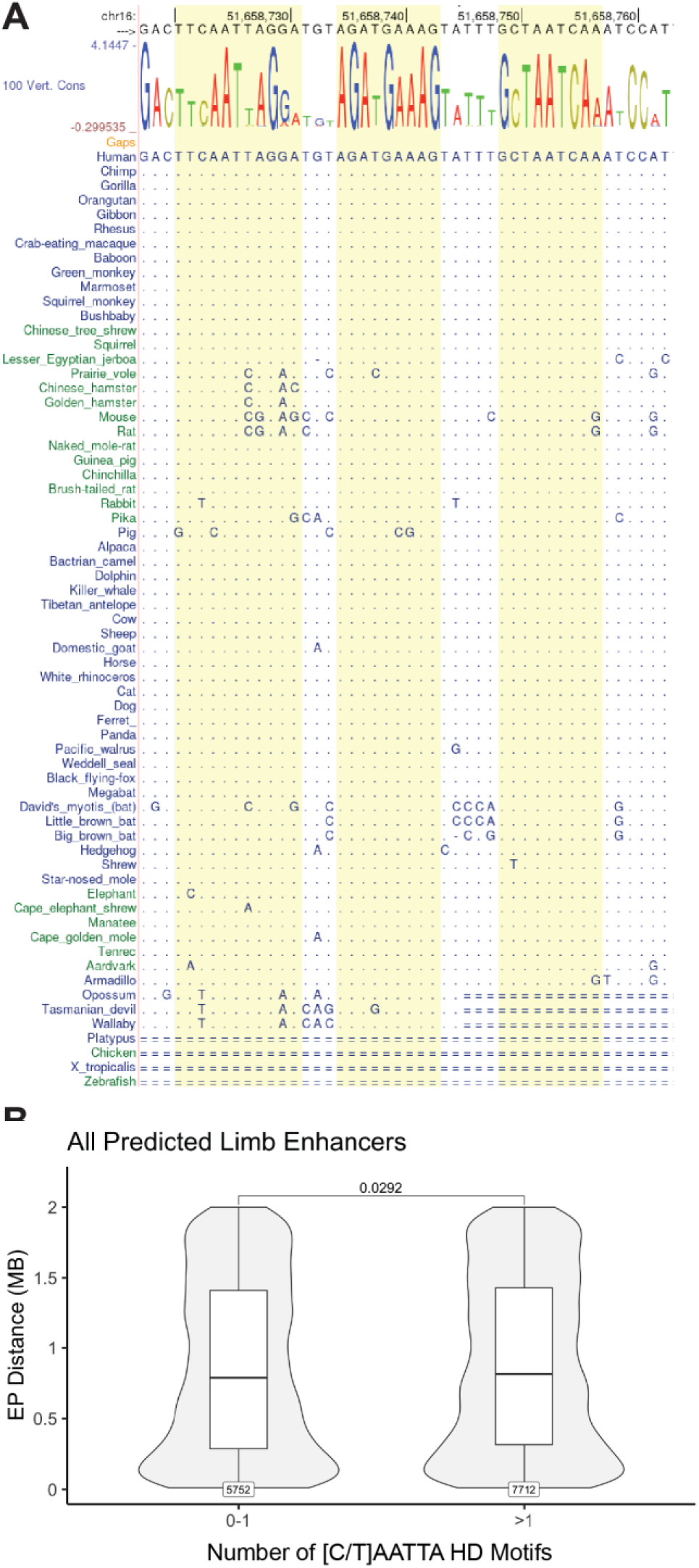
Evolutionary conservation of TF motifs in the REX element. (**A**) Multiple sequence alignment (from UCSC genome browser) across 64 vertebrate species showing conserved LHX2, LEF1, and LHX9 TF motifs in the REX element **(B)** Distribution of enhancer-promoter distances for all predicted limb enhancers with 0-1 compared to more than 1 [C/T]AATTA HD motifs.

**Supplementary Figure 7:**
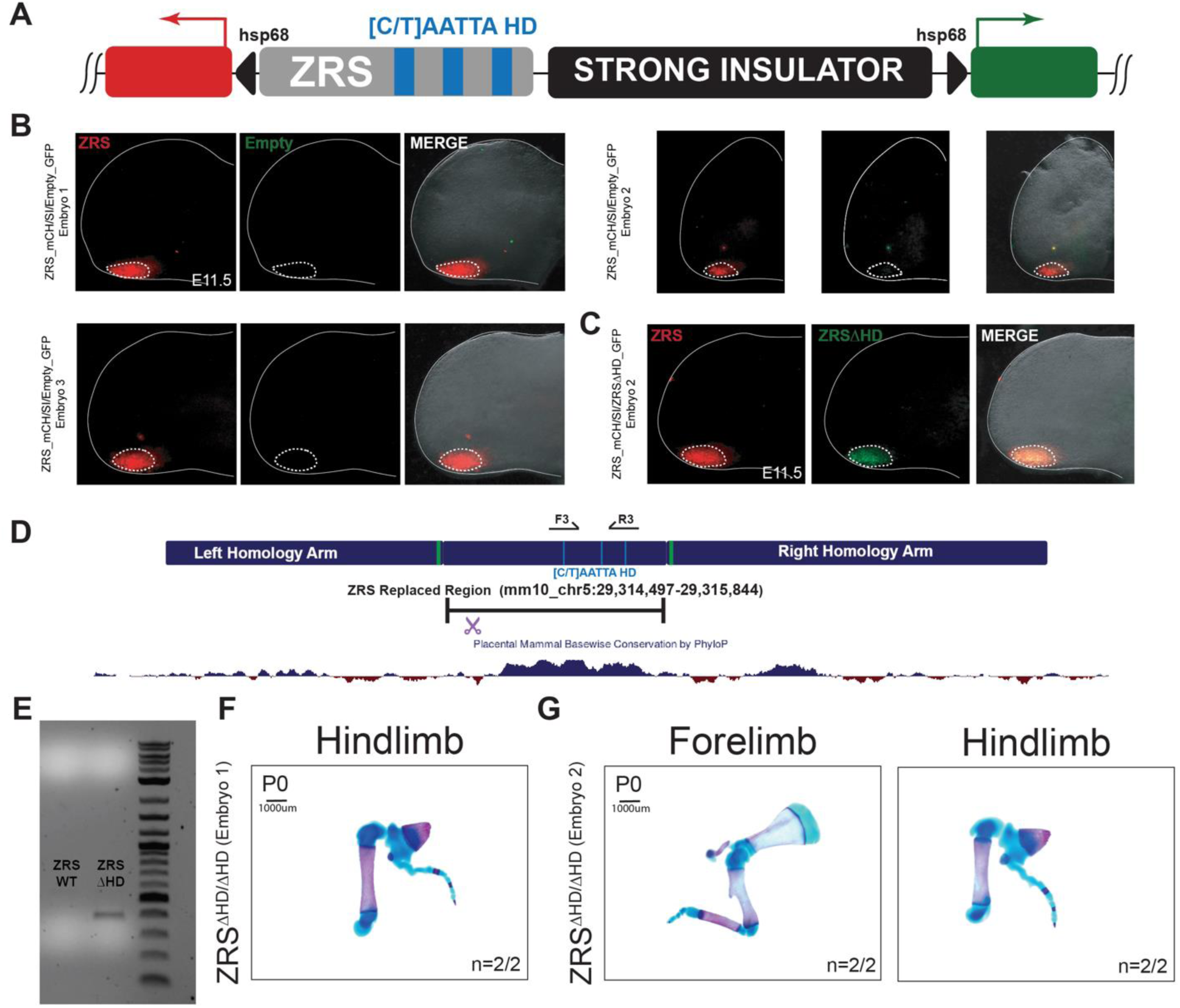
Analysis of [C/T]AATTA sites within the ZRS enhancer. (**A**) Cartoon model of the ZRS-mCH/Empty-GFP dual-enSERT (Hollingsworth et al., 2023) vector (**B-C**) Fluorescent images of E11.5 hindlimbs for mice with a single integration of the ZRS-mCH/Empty-GFP (**B**) ZRS-mCH/ZRSΔHD-GFP (**C**) constructs at H11. Solid outlines the limb bud while dotted lines encircle the region quantified in Fig 6. Lack of GFP fluorescence in Empty_GFP embryos demonstrates that a proximal enhancer is required for activity and that the strong insulator blocks the other enhancer. (**D**) Model detailing the genotyping strategy for ZRSΔHD mice. Primers F3 and R3 are designed to anneal to the mutated [C/T]AATTA HD sites but not the endogenous sequence. (**E**) Example genotyping results demonstrating that ZRSΔHD allele is distinguishable from wild type. (**F-G**) Forelimb and hindlimb images for all the ZRSΔHD/ZRSΔHD mice collected.

**Supplementary Figure 8:**
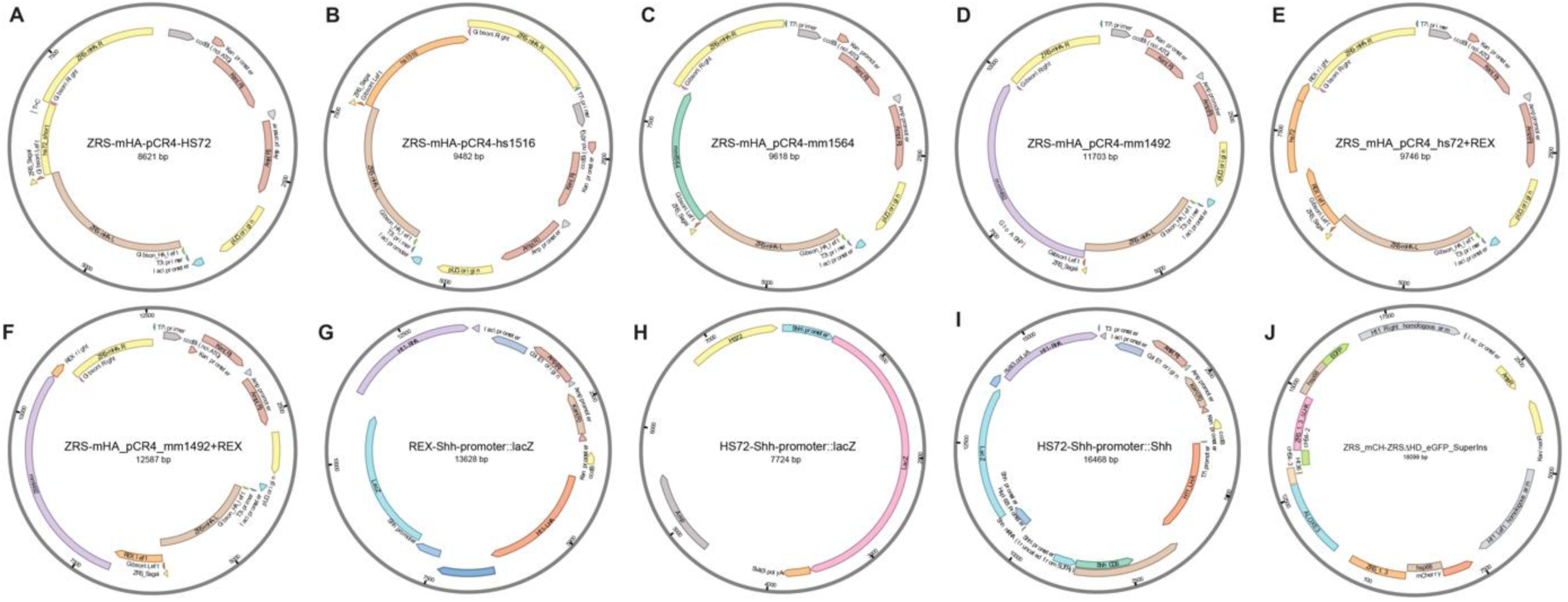
Transgenic constructs used in the study. (**A-F**) Constructs containing the targeting vector that was used in order to replace the ZRS with enhancers of interest. Constructs show in (**G-J**) contain targeting vectors for the H11 locus (Kvon et al., 2020) **(G)** REX-Shh-promoter::lacZ has the regions around the HS72 core enhancer, which contain the REX element (see **Fig S5A**) driving lacZ expression under a minimal *Shh* promoter. **(H)** HS72-Shh-promoter::lacZ contains the HS72 enhancer driving *lacZ* expression under a minimal *Shh* promoter. **(I)** HS72-*Shh*-promoter::*Shh* contains the HS72 enhancer driving *Shh* and lacZ expression under minimal *Shh* promoters. (**J)** ZRS_mCH/ZRSΔHD_eGFP_SuperIns contains the ZRS enhancer driving mCherry and ZRSΔHD enhancer driving GFP separated by a strong insulator, allowing for direct comparison of enhancer activity (Hollingsworth et al., 2023).

## Supplementary Tables

**Table S1**. **Barcode Metadata for Single-Cell Multiomics.** All meta-data outlined for each barcode, including the number of UMI, genes, ATAC fragments, nucleosome signal, transcription start site (TSS) enrichment, MACS2-calculated peaks, and cell type annotation.

**Table S2. Long-Range Motif Enrichment Results.** Motif enrichment for functionally validated (Tab 1) and putative (Tab 2) long-range enhancers. FDR, false-discovery rate; Fold change (Long range / short range motif instance).

